# REAL-TIME AT-LINE MONITORING OF INFLUENZA VIRUS IN CELL CULTURE BY A SURFACE PLASMON RESONANCE BIOSENSOR

**DOI:** 10.1101/2023.03.16.532923

**Authors:** Laurent Durous, Blandine Padey, Aurélien Traversier, Caroline Chupin, Thomas Julien, Loïc J. Blum, Christophe A. Marquette, Manuel Rosa-Calatrava, Emma Petiot

**Affiliations:** Univ Lyon, GEMBAS Team, CNRS UMR 5246, INSA, CPE-Lyon, ICBMS, Université Claude Bernard Lyon 1, Villeurbanne, France; VirPath Team, Centre International de Recherche en Infectiologie (CIRI), Inserm U1111, CNRS UMR5308, Ecole Normale Supérieure de Lyon, Université Claude Bernard Lyon 1, Université de Lyon, Lyon, France; VirNext, Faculté de Médecine RTH Laennec, Université Claude Bernard Lyon 1, Université de Lyon, Lyon, France; CPE Lyon, Bâtiment Hubert Curien, 43 bd du 11 Novembre 1918, Villeurbanne, France

**Keywords:** Influenza virus, On-line sensors, Process Analytical Technology (PAT), Surface Plasmon resonance (SPR), Viral production process

## Abstract

Since the early 2000’, regulation agencies have encouraged viral vaccine manufacturers to implement in-process and real-time monitoring tools in production processes. Even if more assays have been recently developed, none of the novel viral particle quantification technologies can monitor virus levels and their secretion kinetics within production vessels. Vaccine manufacturers still rely on offline cell-based infectivity assays and antigen amount quantification to monitor their processes. The present study describes the development of the first automated biosensor for at-line monitoring of influenza virus production. It involves coupling a fetuin-based SPRi quantitative biosensor with an automated sampler of culture broth and a consecutive clarification setup via an acoustic filter. The SPRi response of different viral strains produced in two distinct cell production platforms was qualified. We demonstrated that fetuin-based quantitative SPRi is a robust, potency-indicating, and universal analytical technology for quantifying bioactive influenza virus particles. It was validated with both purified and complex matrices. Finally, an influenza viral production kinetic was monitored *online* for three days. This novel online tool enabled the access in real-time to total bioactive viral particles from early production phases (8hpi).

## 1. Introduction

Recent years have seen the development of several cell culture-based influenza vaccine processes (Genzel and Reichl, 2009a). They represent strategic alternatives to the historical split-inactivated vaccine produced by ovoculture in case of influenza pandemic outbreaks (Genzel and Reichl, 2009b; Milián and Kamen, 2015; Soema et al., 2015). The most promising strategies for large-scale production rely on suspension cell culture (Gränicher et al., 2019; Lohr et al., 2012; Petiot et al., 2018).

In parallel, FDA’s Process Analytical Technology (PAT) and Quality by Design (QbD) guidance encouraged manufacturers to introduce novel online technologies within their processes to follow both the bioproduct’s critical quality attributes (CQA) and associated the critical process parameters (CPP). Thus, several online probes were implemented to monitor CPP. For example, viable cell concentration is now accessible with capacitance probes (240–243), while cell nutrients and by-products are analyzed thanks to Near-Infrared or Raman Spectroscopy (Petiot et al., 2010; Whelan et al., 2012). More recently, capacitance and fluorescence probes were applied to monitor infected cells, thus unlocking access to the production kinetics of various viruses (Ansorge et al., 2011; Pais et al., 2019; Petiot et al., 2017; Petiot and Kamen, 2012). Although such tools bring new insight into the viral production processes, they do not yet provide information on the viral particle or antigen content. They often require intensive calibration and never give direct access to the bioactivity of the produced virus. Thus, the tedious cell-based infectivity assays remained the routine virus quantification methods for both upstream (USP) and downstream (DSP) viral production processes. Such assays present high variability and a week-long experiment, irrelevant to PAT tools and real-time analysis. Here, biosensors can be tangible assets with their capacity to detect biologically active products. An increasing number of scientific work report SPR-based quantitative assays applied to the analysis of a wide variety of bioactive molecules and applications in the bioprocess monitoring field (Dorion-thibaudeau et al., 2017; Karlsson, 2016; Karlsson et al., 2018; Pol et al., 2016). Interestingly, in the last ten years, SPR-based biosensors were applied to detect and quantify a wide range of viruses, including influenza strains (Durous et al., 2019c; Khurana et al., 2014; Nilsson et al., 2008; Nilsson et al., 2010), SARS-COV2 (Shang et al., 2020), enterovirus, norovirus (Singh, 2017), African swine fever virus (Capo et al., 2022), etc....But the implementation of this technology for bioprocess monitoring is relatively slow. A unique work presents at-line Monoclonal antibodies monitoring during a CHO culture process by Surface Plasmon Resonance (SPR) (Dorion-thibaudeau et al., 2017). In the virus bioproduction field, the community evaluated several physical quantification techniques to reach in-process monitoring of the viral particle (e.g., TRPS, NTA, FFF-MALS, flow virometry) (Faez et al., 2015; Makra et al., 2015; Rossi et al., 2015; Zhou et al., 2011). They are all considered at-line tools that permit particle counting less than an hour after sampling (Heider and Metzner, 2014), but none were installed online.

In previous work, we presented a fetuin-based SPRi assay to quantify influenza vaccines and virions. This study demonstrated similar or improved quantification performance to standard SRID or infectivity assays (Durous et al., 2019b). We now aim to show the broad applicability of this tool by first generalizing this analytical approach for multiple production strategies (egg, adherent MDCK, suspension avian DuckCelt®-T17) and sample qualities (several influenza strains, different sample purification stages). Then, we integrated this tool within an automated sampling loop, including a dedicated sample pre-treatment line, to demonstrate the feasibility of real-time monitoring of bioactive influenza production kinetics.

## 2. Material and methods

### 2.1. Reagents

Fetuin from fetal bovine serum, bovine serum albumin, ethyl dimethyl carbodiimide (EDC), N-hydroxysuccinimide (NHS), sodium dodecyl sulfate (SDS), polysorbate 20, trypsin acetylated from bovine pancreas was purchased from Sigma (Saint Quentin Fallavier, France). Sodium acetate buffer and ethanolamine-HCl from Brucker Daltonics (Hamburg, Germany). Phosphate Buffer Saline (PBS) tablets, Pluronic F-68 (P1300), and trypsin (T6763) were acquired from Merck (Darmstadt, Germany). NuPAGE™ LDS buffer was from Thermofischer. All solutions were prepared using Milli-Q water. Oseltamivir carboxylate was purchased from Roche (Bale, Switzerland). Madin–Darby canine kidney cells (MDCK, ATCC CCL34), serum-free Ultra-MDCK, EMEM, and Optipro SFM cell culture media, L-glutamine and penicillin/streptomycin solutions were obtained from Lonza (Amboise, France). Avian DuckCelt^®^-T17 cell lines (ECACC 09071703) were acquired from Transgene. SRID reference antigens were acquired from the National Institute for Biological Standards and Control (NIBSC).

### 2.2. Influenza virus production

Two influenza A subtypes were produced on suspension DuckCelt®-T17 cells. First, the A/Reassortant/NYMC X-179 (H1N1) which is a reassortant prepared by New York Medical College using classical reassortant methodology from A/California/7/2009 (H1N1) virus and NYMC X-157 virus, with the HA, NA and PB1 genes donated from A/California/7/2009 (H1N1) and the other internal genes donated from A/PR/8/34 (H1N1). Second, the A/Panama/07/99 (H3N2), a viral strain part of the vaccine composition of 2003-2004. DuckCelt®-T17 cells were grown in suspension, and Optipro SFM medium supplemented with L-glutamine (2 mM), penicillin (100 U/mL), streptomycin (100 U/mL), and 0.2% (w/v) Pluronic F-68. Before virus production, DuckCelt®-T17 cells were seeded at 0.7.10^6^ cells/mL and amplified to reach between 0.8.10^6^ and 1.4.10^6^ cells/ml before infection. Infection was performed at a multiplicity of infection (MOI) of 10^-3^ virus/cell in the presence of 0.5 μg/mL trypsin. Productions for offline evaluation were run in 500 mL shaker flasks at 37 °C in a CO_2_ Khüner incubator (ISF1-X, Khüner) with an agitation speed of 120 rpm. *Online* experiments were achieved thanks to the connection of the SPRi equipment to a 500 mL Bellco spinner flask in the CO_2_ incubator (37°C, 5% CO_2_, 120 RPM). Cell growth and viability were monitored every two hours through manual sterile sampling. For biosensor calibration, assays were performed from A/Reassortant/NYMC X-179 (H1N1) influenza virus produced on adherent MDCK cells, as in our previous study. MDCK cells were grown in 175 cm^2^ T-flasks in Ultra-MDCK serum-free medium supplemented with L-glutamine (2 mM), penicillin, and streptomycin (100 U/mL). At 70-80% of confluence, influenza viruses were inoculated at a multiplicity of infection (MOI) of 10^-3^ viruses/cells. Cell cultures were collected daily and clarified using 2000g centrifugation.

### 2.3. SPRi online-line setup

#### 2.3.1. Automated sampling and pre-treatment line

An automated sampling and pre-treatment line set up was built (**FIG 1**). A Biosep 10L acoustic filter (Applikon Inc., Foster City, CA) related to the bioreactor thanks to two peristaltic pumps (P-1, GE Healthcare), one for medium recirculation and one for sample withdrawal (Henry et al., 2004). The recirculation flow rate was settled to 6 mL/min while the harvest pump flow rate was 2 mL/min. The sampling frequency was set to fill an HPLC valve loop with 500 μL. Thanks to a microcontroller board, the HPLC valve was commanded (**Supplementary Information 4**). The SPRi syringe pump was used to realize sample diafiltration by suction/discharge before filling dedicated vials within the SPR device autosampler.

**Figure 1.**
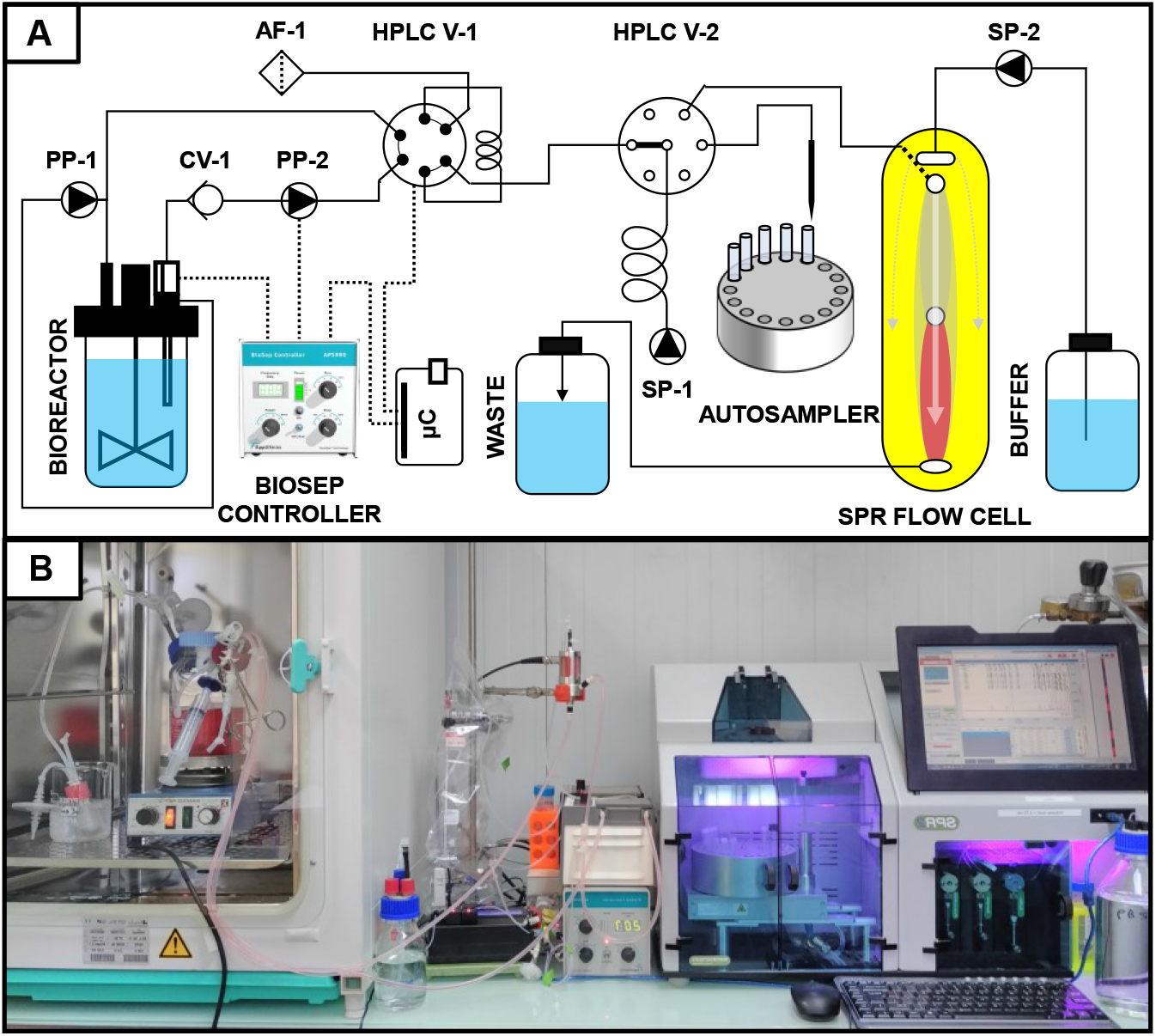
Graphical Abstract. Automated online sample treatment and analysis setup. A_Process and Instrumentation Diagram. AF: Air filter; PP: Peristaltic pump; CV: Check valve; SP: Syringe pump; μC: microcontroller. B_Image of the entire online experimental setup.

#### 2.3.2. SPRi quantitative assay and bioactive virus concentration determination

The quantitative assay was performed by Imaging Surface Plasmon Resonance (SPRi) using the SPR-2 instrument from Sierra Sensors, as described in Durous et al. 2019 (Durous et al., 2019b). Both sensing and reference surfaces were functionalized with immobilized fetuin (200 μg/mL, pH=4.5). Then, the reference surface was treated with bacterial neuraminidase (2.5 U/mL) to cleave α-2,3, α-2,6, or α-2,8 terminal sialic acid. Clarified cell culture samples were collected every hour, thanks to the automated sampling and pre-treatment line, and were injected into the SPRi autosampler. SPRi autosampler vials were added with oseltamivir carboxylate final concentration of 10 μM. 80 μL samples were injected in duplicates into the SPRi flow cell at a 40 μL/min flow rate, leading to a 2 min interaction time. The running buffer was composed of PBS with 0.05% (v/v) of Tween 20 and 10 μM oseltamivir carboxylate. Surface regeneration was performed by injecting 25 μL of PBS buffer containing 0.25% (v/v) of SDS. Initial binding rates [RU/s] were calculated by linear regression of the response during the whole association phase, corresponding to mass transport limitation (MTL) conditions (Karlsson, 2016). A moving average filter (n=3) was applied for data smoothing on online production kinetics.

### 2.4. Infectivity assays: Infectious viral particles (IVP) quantification

Infectious influenza samples (IVP/mL) were titered by plaque forming units (PFU) assays (Dulbecco, 1954). Infection of confluent MDCK cells cultivated with serial dilutions of the virus was performed in 6-well plates and EMEM medium. 800 μl of viral suspensions were deposited on the cell monolayer and incubated for 1 h at 37 °C under continuous shaking. MDCK cells were washed with EMEM medium and overlaid with Nobel agar (1.1%) mixed 1:1 with modified MEM composed of 2x MEM, 200 U/mL of penicillin-streptomycin, and 2 μg/mL of trypsin. Infected cell cultures were incubated for 3 days at 37 °C, 5% CO_2_, before visual inspection of the cytopathic effect.

### 2.5. Total particle (TP) determination by TRPS

Total particles were quantified by the Tunable resistive pulse sensing technique (TRPS) using qNano Gold (IZON Science, Lyon, France). Samples were diluted at 1/5 in PBS buffer before analysis. Analyses were performed using thermoplastic polyurethane membranes with a size-tunable nanopore of 150 nm (NP150). According to the manufacturer’s instructions, calibration was performed using carboxylated polystyrene beads standards (CPC150, Izon Science Ltd., Lyon, France).

### 2.6. Quantification of Hemagglutinin (HA) antigen

#### 2.6.1. Hemagglutination assay for potency evaluation

As described, the HA titers of clarified and heat-denatured supernatant samples were quantified by hemagglutination assay in v-bottom 96-well plates (Eisfeld et al., 2014). Aliquots of 50 μl were serially diluted 1:2 in PBS. They were further incubated for 1 hour at room temperature with 50 μl of 0.5% chicken red blood cells (RBCs). Then, HA titers were evaluated as follows, the last dilution presenting hemagglutination was transposed as the HA units/50μl. Relative HA content presented in the graphs was assessed as the ratio of HA titers and expressed in %.

#### 2.6.2. HA protein content by Western Blot (WB) densitometry

HA protein content was analyzed thanks to quantitative western blot densitometry. Clarified viral production supernatants were incubated with LDS lysis buffer supplemented with dithiothreitol 0.1 M and heat-denatured for 5 min at 95 °C. Samples were then loaded onto a 10% SDS-PAGE gel for migration and later transferred to a nitrocellulose membrane. Western blot analysis was performed using a universal anti-hemagglutinin antibody cocktail (F211-10A9 and F211-11H12, 6 μg/mL) as primary antibodies set provided by NRC-Montreal laboratory incubated overnight at 4°C (Manceur et al., 2017). Anti-mouse HRP secondary antibody (1/5000 dilution) was incubated for 1 hour at room temperature. The densitometry of WB bands was analyzed thanks to the Image Studio software (LICOR Biosciences, Lincoln, NE).

## 3. Results and discussion

### 3.1. Mass-transport limitation evaluation for bioactive influenza virus particles

Our fetuin-based SPRi biosensor operates under full mass-transport limitation conditions for determining bioactive virus concentration. These conditions were confirmed thanks to the serial dilution of H1N1 influenza virus samples produced in MDCK cell culture (1.35 x 10^8^ TCID50.mL^-1^) and the processing of samples at four different flow rates (5, 10, 20, and 40 μL.min^-1^). For SPRi biosensors operating under mass-transport limitation (MTL) conditions, the binding rate (dR/dt) shall be directly correlated to the analyte bulk concentration according to the following equation (Karlsson, 2016; Pol et al., 2016) :

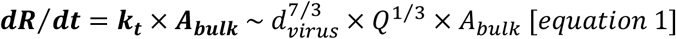

Where ***k_t_*** [RU.s^-1^.VP^-1^.mL] is the mass transport coefficient, ***A_bulk_*** [VP.mL^-1^] corresponds to analyte bulk concentration, ***Q*** is the operating flow rate, and ***d_virus_*** is the virus diameter (see equation simplification in **Supplementary information 1**). The binding rate of influenza particles is linearly correlated with total virus particle content in mass-transport limitation conditions (Durous et al., 2019b). The slopes of the curves obtained for the binding rate in this experiment allow us to determine the mass transport coefficient, *kt,* at different flow rates (**FIG 2.A)**. The r^2^ coefficient obtained for the correlation between *k_t_* values and Q^1/3^ at different flow rates confirms strong linearity (r^2^=0.99; **FIG 2. B**). It validates the hypothesis of transport limitation for influenza virus binding to the SPRi surface (see details in **Supplementary information 1**).

**Figure 2.**
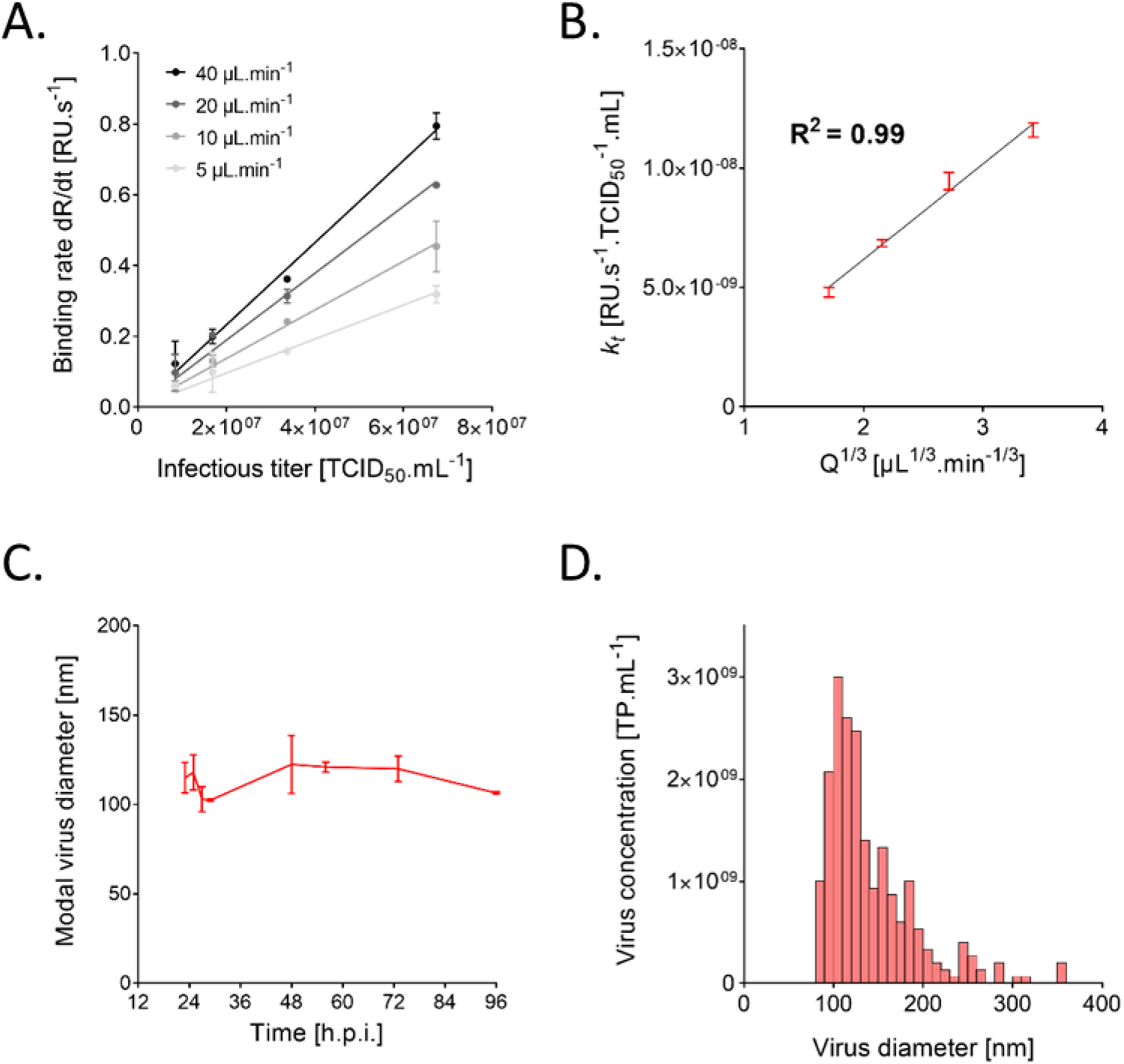
Evaluation of mass-transport limitation for H1N1 A/California/07/09 influenza virus produced in MDCK cell culture. A_Calibration curves obtained at 4 flow rates. B_Linear regression between the corresponding mass-transport coefficients evaluated at 4 flow rates and Q ^1/3^. C_Kinetics of the modal diameter variations of the H1N1 virus particles population derived from MDCK cell culture according to the time post-infection evaluated by TRPS (n=2). D_Size distribution of H1N1 virus particles derived from MDCK cell culture evaluated by TRPS (48 hpi).

While viruses are large macromolecular entities that diffuse slowly (*D* ~ 5.10^-12^ m^2^.s^-1^) (Parupudi et al., 2017), which makes them particularly suitable for such SPRi quantitative analysis, their diameter variation is also well known (Moulès et al., 2011). Consequently, the size of the virion all along the bioproduction had to be evaluated since it could impact the mass transport coefficient and the robustness of our assay.

The size of the H1N1 influenza virus population produced on MDCK cells culture was measured throughout 96 hours of production by tunable resistive pulse sensing (TRPS) analysis. The influenza virus population was shown to be monodisperse with a quasi-Gaussian diameter distribution around 114 ± 8.3 nm (RSD = 7.3%) (**FIG 2. C**). A negligible evolution of the virus particle diameter during the production process was then demonstrated (**FIG 2.D**).

Assays were also performed to validate that our SPRi biosensor can selectively detect bioactive virus particles (i.e., carrying bioactive HA antigens). As depicted in **FIG3,** the SPRi assay was used to characterize heat-treated H3N2 virus degradation after increasing inactivation times. The biosensor was able to follow a time-dependent decrease in the virus bioactivity measured by standard potency assays (infectivity (PFU) and hemagglutination assay (HA))(**details in Supplementary Information 3)**.

**Figure 3.**
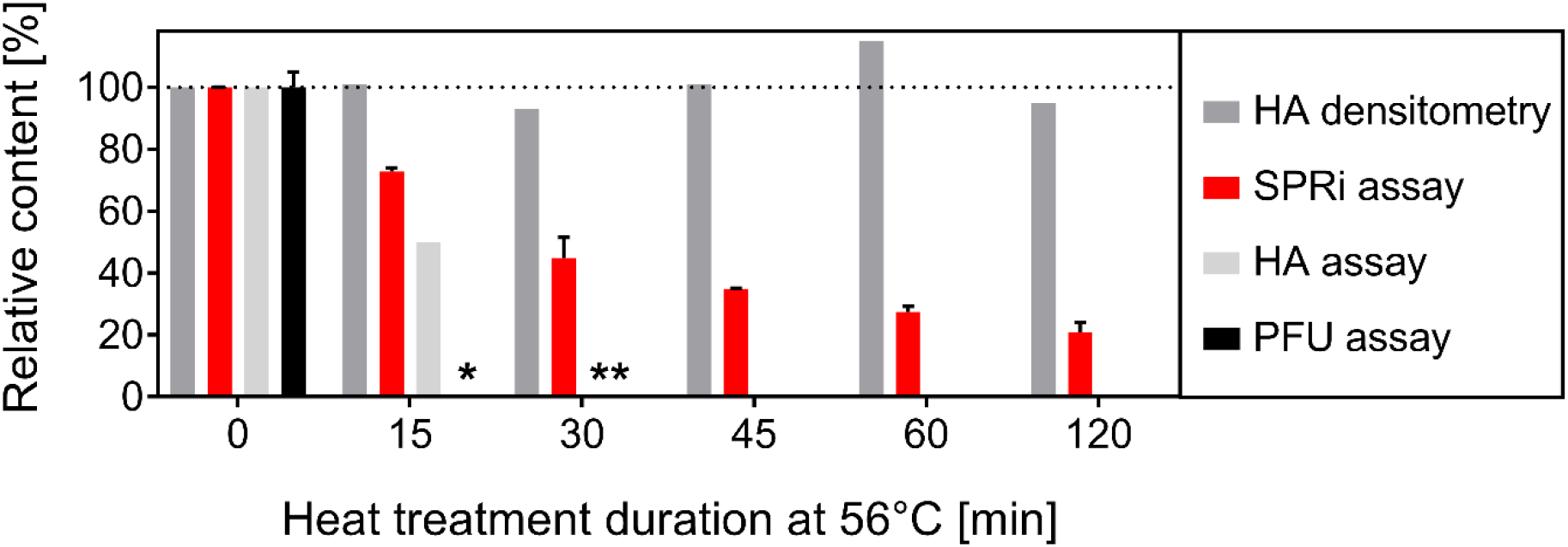
Thermic degradation of H3N2 A/Panama/07/99 influenza virus following various incubation times, determined by HA densitometry (n=1), SPRi analysis (n=4), HA assay (n=2) and infectivity assay (n=2). *Infectivity titer <10 PFU.ml-1,** HA titer < 2 HAU. (Detailed data in Supplementary Information 2)

### 3.2. Offline monitoring of bioactive influenza production kinetics

SPRi was applied to monitor influenza production kinetics in avian suspension cells (Duckcell-T17) cultivated in 200 mL shake flasks. This production strategy was chosen to consolidate the results obtained in previous work on MDCK adherent cells (Durous et al., 2019a). It aims to confirm the applicability of this quantification strategy for novel suspension cell-based productions currently under development (Gränicher et al., 2019; Lohr et al., 2012; Petiot et al., 2018). The strategy was deployed for a novel influenza strain, the A/Panama/07/99 (H3N2) influenza strain.

Characteristic production profiles were observed for H3N2 strains (Petiot et al., 2011a; Petiot et al., 2015; Petiot et al., 2018). They have been characterized thanks to infectivity assay (PFU) and cell growth/ death monitoring (see **FIG 4**). After one day of cell growth to reach a target cell concentration at infection (CCI) of 0.5 to 2 x 10^6^ cells/ml, cells were infected at a low multiplicity of infection (MOI: 10^-3^). Thus, after 32 hours post-infection (hpi), cell death appears with a clear drop in cell viability and a loss of total viable cells. This trend goes with the infective virus particle secretion, which reaches 2.6 x 10^6^ PFU/ml. Production trends and maximal levels (10^6^ pfu/ml) are similar to the titers previously published with Duckcellt-T17^TM^ cell line or other suspension cell platforms (Lohr et al., 2010a; Lohr et al., 2012; Petiot et al., 2011b). The production levels were monitored in parallel using the initial binding rate of SPRi signals, as presented in **FIG 4.A & B**. The binding rate trends are observed in **FIG 4.A** demonstrate a progressive increase. It corresponds to an increased interaction of the fetuin-modified surface with viral particle hemagglutinins and indicates the increase of total bioactive influenza virus particle presence. Indeed, the offline correlation between SPRi binding rate and the total influenza virus particles quantified by TRPS proved reliable with R^2^ >0.80 (**FIG 4.B**). On the contrary, the influenza infectious titers trends demonstrate a maximal titer at 32 hpi. It evidences the evolution, over the process phases, of the proportion of infectious viral particles among the total bioactive viral particles carrying hemagglutinin. Such behavior has already been described in several cell-based production studies. It can correspond to either the progressive degradation of infective virus particles over time (Durous et al., 2019b; Genzel et al., 2014; Lohr et al., 2010b; Petiot et al., 2011b; Petiot et al., 2018; Transfiguracion et al., 2015), or the cellular misconception of the viral particles. Consequently, SPRi biosensors allow more reliable monitoring of the total bioactive particles while being more accurate than infectivity assays for counting particles carrying hemagglutinin. Also, the standard sampling rate in viral production processes is once a day (Lohr et al., 2009; Lohr et al., 2010b; Petiot et al., 2011b; Petiot et al., 2018), SPRi analyses have the added value to reach a high-frequency quantification rate of one sample per hours. Consequently, in-process real-time monitoring of bioactive influenza viruses can be envisaged. It is proved in the next section by implementing SPRi as an online monitoring strategy.

**Figure 4.**
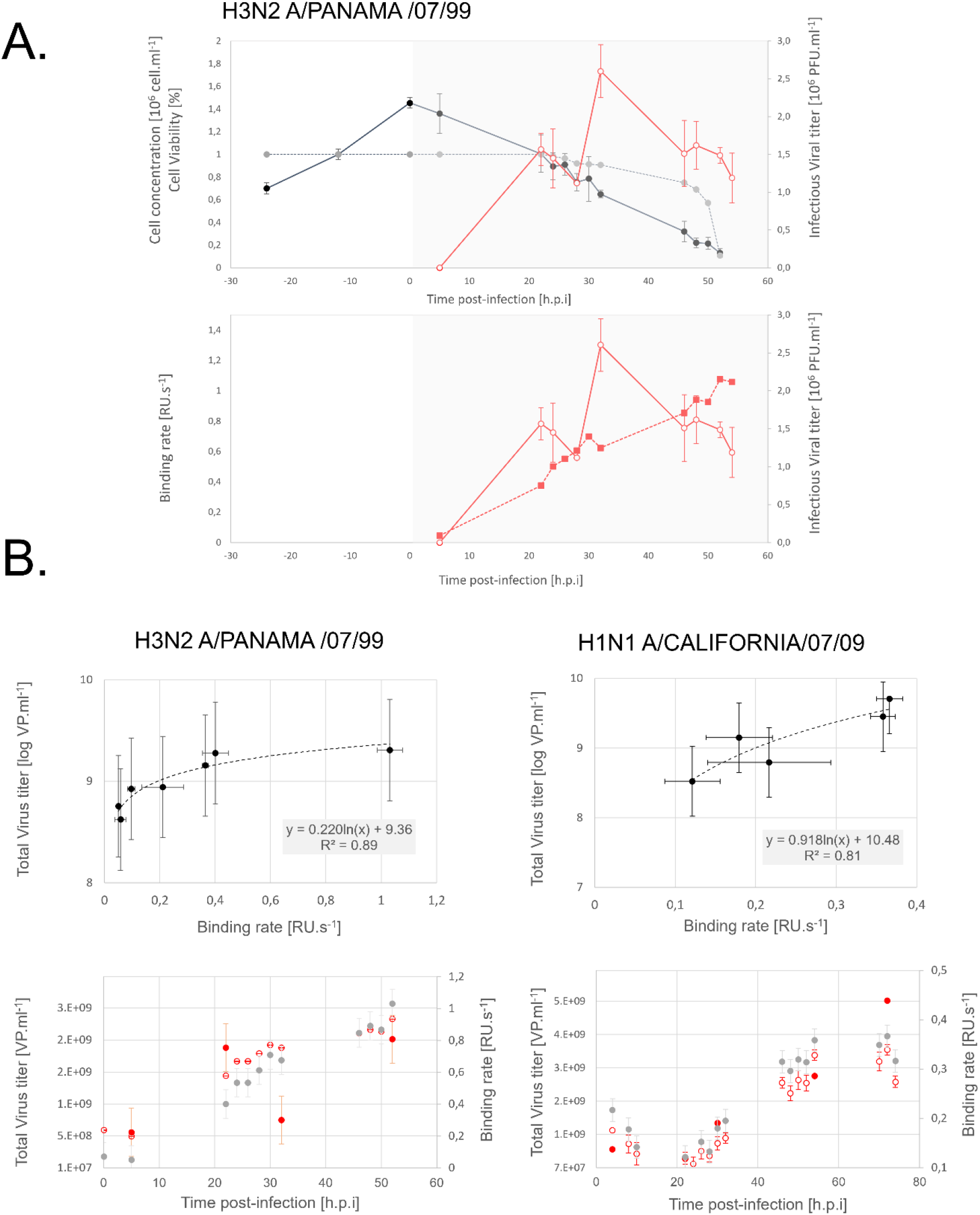
Offline monitoring of A/Panama/07/99 (H3N2) and A/reassortant/NYMC X-179 (H1N1) influenza strains by SPRi biosensor. A_Example of A/Panama/07/99 (H3N2) production kinetic (production condition: MOI 0.01; trypsin 1μg.ml^-1^): Viable cell concentration (●); Cell viability 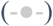; infectious virus particle titers measure by PFU 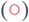; SPRi biosensor binding rate 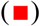. B_Correlation between total virus particles quantified by TRPS and SPRi binding rate (∎). C_Viral production kinetics monitoring: Total virus titer measured by TRPS 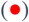; Predicted total viral titer 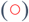; Binding rate trends 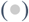.

### 3.3. Online monitoring of bioactive influenza virus during production processes in suspended cell culture

A novel setup integrating an automated sampling and pre-treatment line was built to evaluate the feasibility of real-time monitoring of the viral kinetic using the developed SPRi biosensor (see **FIG 1 Graphical abstract**). The complete design, list of equipment pieces, and microcontroller script are detailed in **Supplementary information 5**.

The developed biosensor was implemented online to monitor the production of bioactive virus particles by measuring the initial binding rate at the fetuin-modified SPRi surface. Online automatic sampling of cell culture was performed every hour. The targeted production was an H3N2 virus produced at an MOI of 0.001 virus/cell and from cell culture infected at 0.8 x 10^6^ cell/ml. DuckCell-T17^TM^ was grown in a 500 mL spinner flask installed in a standard CO_2_ incubator. The culture was sampled hourly and clarified thanks to a Biosep acoustic filter. Following clarification, samples were directly injected into SPRi autosampler.

As a first validation step, the auto-sampler and Biosep clarification protocols were compared with manual sampling and standard centrifugation protocols from the same production batch to control the impact of these practices on virus bioactivity. A correlation between the response obtained with both protocols is presented in **FIG 5,** confirming that auto-sampling and clarification lines did not impact the viruses’ binding capacity (**FIG 5.A**: R^2^=0.92). It validates the setup for online influenza virus kinetic monitoring.

**Figure 5.**
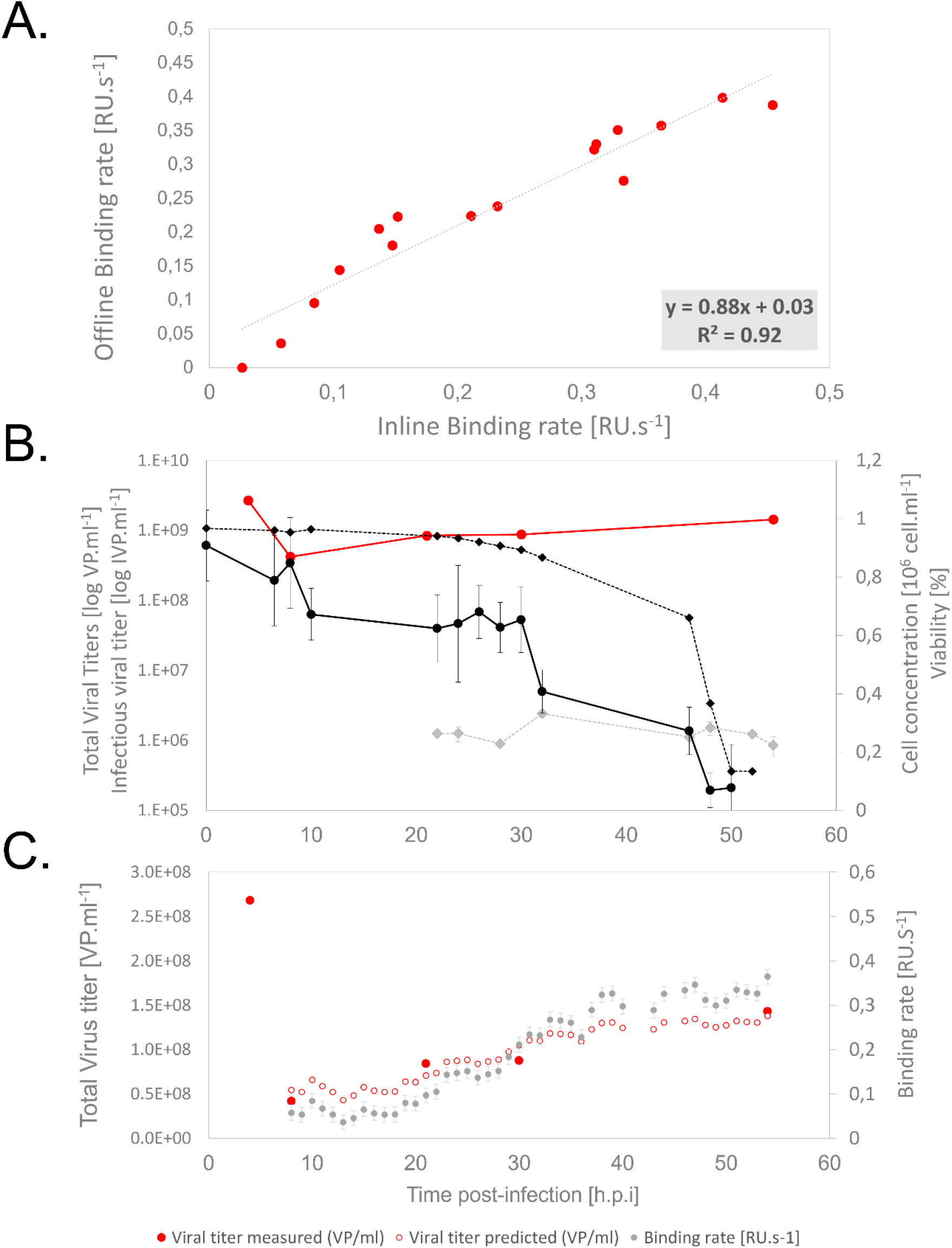
Online monitoring of influenza A/Panama/07/99 (H3N2) virus (MOI = 10^-3^ virus/cell) produced on suspension cell culture (n=2 successive measurements). A_Correlation between at-line and offline SPRi binding rates. B_Virus production kinetics: Viable cell concentration (●), Cell viability (♦); Total virus particles measured by TRPS 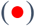 and infectious virus particle titers measure by PFU 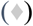. C_ Comparative trends between online viral titers measured by SPRi and reference TRPS: Total virus titer measured by TRPS 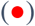; Predicted total viral titer 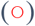; Binding rate trends 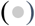.

The monitored H3N2 production kinetic is presented in **FIG 5. B**. Production kinetics is similar to the ones observed in shake flasks (section 3.2). Cell growth stops after infection, and a cell death appears 24 hours post-infection, confirming the infection process is ongoing with a concomitant drop in viable cell number. A significant binding rate was recorded as early as 8 hpi with a 0.057 RU.s^-1^. According to previous correlations, this corresponds to a total viral particle titer of 5.43 10^8^ VP/ml. Then, the binding rate was monitored hourly, informing upon the viral release trends over the 56 hours of the viral production process. Thus, several successive increases in the binding rate were observed (see arrows **FIG 5C**) at 10 hpi, 23 hpi, 29 hpi, and 37 hpi before stabilizing. As described by previously published work, the such viral release could correspond to infection cycles occurring in cell-based processes optimized at low MOI. Indeed, influenza infection cycles were defined to commonly last about 8 to 10 hours (Petiot and Kamen, 2012). We could then hypothesize that at least four viral cycles shall be observed for the present influenza H3N2 A/Panama/07/99 production process (0-10 hpi / 10-24 hpi / 24-30 hpi/ 30-37 hpi).

Thus, the developed biosensor enabled access to bioactive influenza virus production kinetic hourly and early production phase monitoring for the first time. This setup represents a unique and first available technology for real-time tracking of bioactive influenza virus particles and production processes. It was demonstrated to apply to both adherent and suspension cell cultures. Also, an apparent discrepancy between the binding rate and standard infectivity assay indicated that total bioactive particle count should be a more indicative parameter for process and productivity monitoring.

## 4. Conclusion

A fully automated SPRi assay was developed to monitor bioactive influenza virus particles. This strategy was deployed on suspension cell culture bioprocess but was quickly and easily transferable to adherent cell-based systems. We proved that, unlike immunoassays, the SPRi response does not depend on the virus strain or cell expression platforms. Glycan-based quantitative SPRi can rapidly, precisely, and universally quantify influenza bioactive virus particles while not necessitating specific reagents. Such proof-of-concept bridges the gap toward developing biosensors for online cell culture monitoring. It also confirms that bioactive total viral particles in cell culture are an additional critical quality attribute to focus on while developing bioproduction processes. Combining SPRi monitoring with recent orthogonal techniques evaluating real particles (e.g., TRPS, flow virometry...) is a powerful approach to better understanding viral secretion dynamics. In that sense, such a tool could strongly support PAT and QbD strategies introduced by the FDA. It is also a strategic solution to solve the quantification problems of virus-like particles (VLP), which are non-infectious but still present bioactive antigens at their surfaces.

## Supporting information

Supplementary Informations

## Acknowledgments & Competitive interests’ disclosure

L. Durous acknowledges the Auvergne Rhône-Alpes Region for his Ph.D. grant (ARC 1 Santé). The authors would like Brucker Daltonics SPR (Sven Malik & Klaus Wiehler) team for their support regarding the SPR-2/4 apparatus and software. We thank Aziza P. Manceur and Amine Kamen for providing universal anti-hemagglutinin antibodies.

## SUPPLEMENTARY INFORMATION

### SUPPLEMENTARY INFORMATION 1 Impact of virus size variation on the mass transport coefficient *k_t_*: Schematic representation of the transport of virus particles in the biosensor flow cell

**SUPPLEMENTARY FIG 1.**
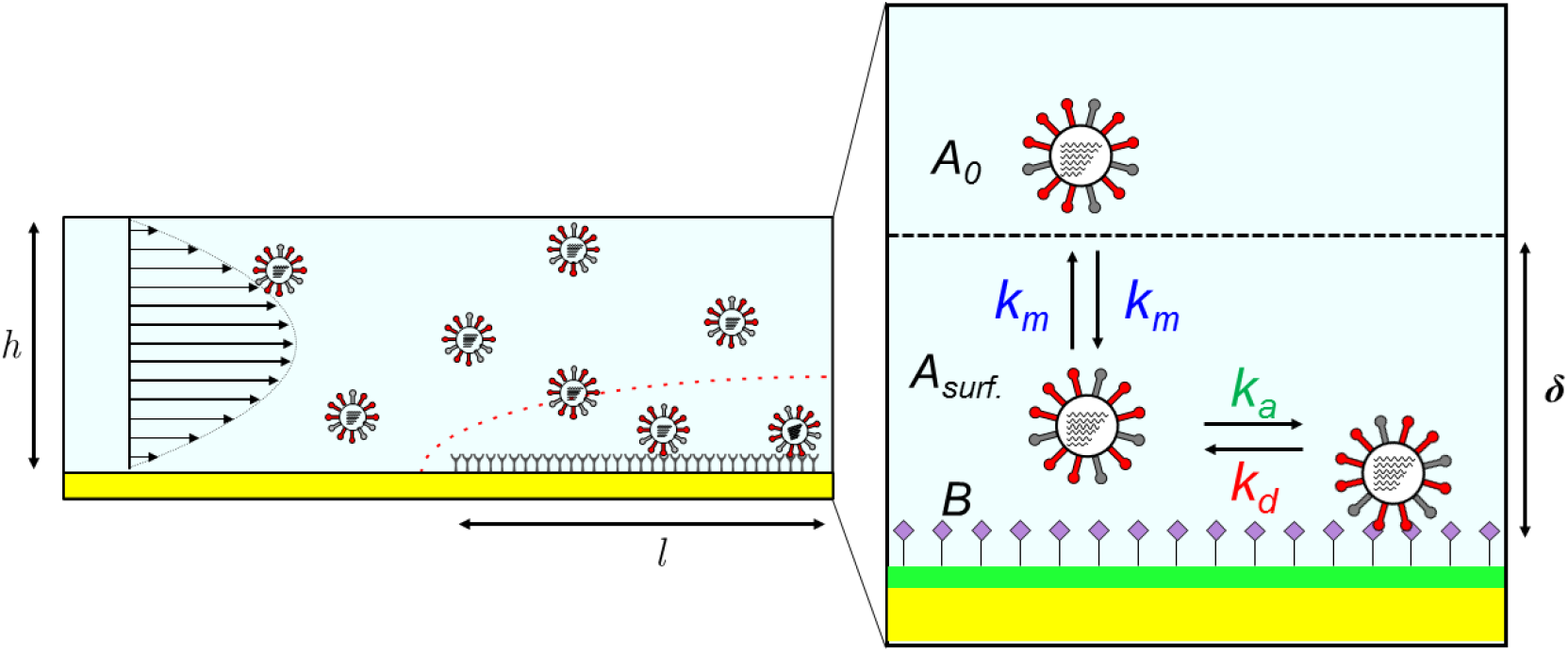
Impact of virus size variation on the mass transport coefficient *k_t_*: Schematic representation of the transport of virus particles in the biosensor flow cell.

The diffusive flux *J* [*mol. m*^2^. *s*^-1^] of an analyte through the depletion zone (*δ*) of a biosensor is given by (Sjölander and Urbaniczky, 1991; Squires et al., 2008):

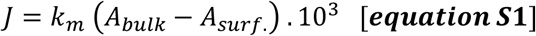

Where *k_m_* [*m. s*^-1^] is the heterogeneous mass transport coefficient and *A_bulk_* and *A_surf_*. [*mol. L*^-1^] correspond respectively to the analyte concentrations in the solution and close to the sensor’s surface. Factor 10^3^ is used as a conversion factor: *mol. m*^-3^ ↔ *mol. L*^-1^. For a rectangular flow cell with one sensing wall, the heterogeneous mass transport coefficient has been analytically derived (Karlsson, 2016; Pol et al., 2016) :

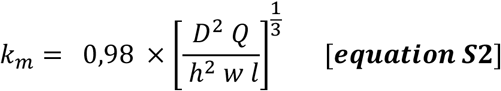

Where *D* [m^2^.s] is the diffusion coefficient of the analyte, Q [m^3^.s^-1^] is the volumetric flow rate, and h, w, and l [m] respectively correspond to the height, the width and the length of the flow cell. Surface-based biosensors can be operated in conditions of mass transport limitation, where the binding rate of biomolecules to the biosensor’s surface is limited by the diffusion phenomenon, predominating over the association phenomenon. Under these conditions, the bulk concentration of the analyte is proportional to the initial binding rate dΓ/dt according to the relation:

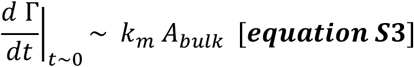

Γ [*mol. m*^2^] being the surface-bound density of the analyte. The corresponding response of SPR biosensors (e.g., the initial binding rate in terms of response unit) also takes other parameters into account:

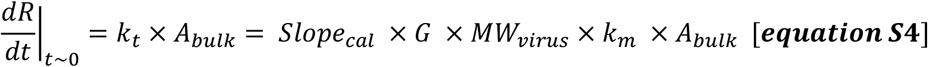

Slopecal is a calibration coefficient between the SPR response and the local refractive index variation, G is a conversion factor between the refractive index variation and the mass surface density, and *MW*_virus_ is the molecular weight of Influenza virus particle (Pol et al., 2016).

Besides its flow rate dependence (*k_t_* ~ Q^1/3^), *k_t_* depends on constants (i.e. geometric flow cell parameters and conversion factors) and analyte-related parameters (*D_virus_* and *MW_virus_*).

The size-dependence of the diffusion coefficient of the virus can be evaluated according to the Stokes-Einstein equation:

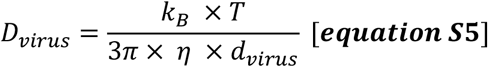

Where *k_B_* represents the Boltzmann constant, *T* the temperature and *η* the viscosity of the medium. The flow cell temperature was kept constant during all experiments. Buffer viscosity was also considered constant as all samples were diluted in the same buffer for calibrations (PBST added with 10μM oseltamivir carboxylate). Additionally, the viscosity of the cell culture medium (Optipro®) was considered identical to PBST buffer, as no serum was added.

While for the molecular weight of influenza virus:

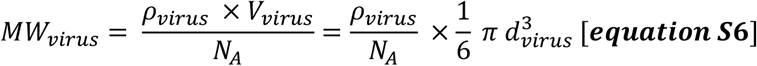

Where *ρ_virus_* represents the density and *V_virus_* the volume of the virus particle and

By coupling **equations S4, S5 and S6**, we can deduce the mass transport coefficient is proportional to the virus particle diameter according to 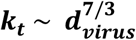.

### SUPPLEMENTARY INFORMATION 2 Qualification of the SPRi biosensor and selection of mass transport limitation conditions for the quantitative analysis of influenza virus particles

#### Calibration of the SPR-2 apparatus

was realized by analyzing glycerol solutions diluted in PBST buffer onto the functionalized sensor. The solutions were of increasing refractive index ranging from 0 to 0.0015 [ΔRIU]. Considering a 1% w/w glycerol solution yields a refractive index increment Δn_s_ of 0.00113 Refractive Index Unit [RIU], we evaluated the system response as:

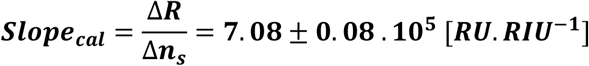

This value is close to the conventional value of **1.0.10^6^ [*RU.RIU*^-1^]** reported in the literature for commercial SPR systems (Tudos and Schasfoort, 2008). The baseline noise of the response was evaluated as σ (R) = 0.1 [RU]. The resolution of the system can be evaluated by the limit of detection S = 3.3* σ / Slope_cal_ = 5.10^-7^ [RIU].

**Supplementary FIG3:**
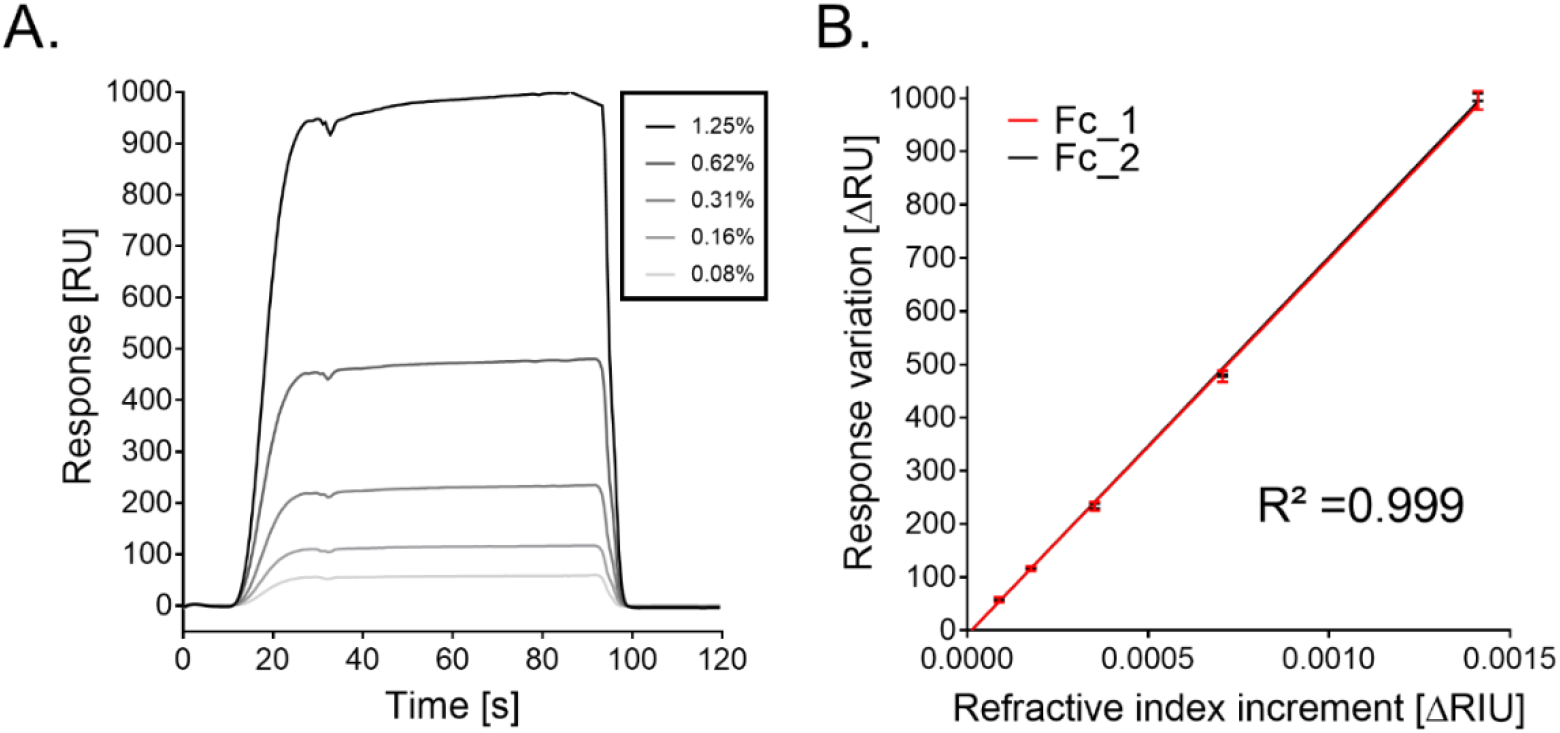
Calibration of the SPR-2 apparatus. A_Sensorgrams of the response for glycerol solutions of increasing concentrations in PBST buffer. B_Calibration curves obtained by difference between response of the reference surface and the sensing surface at the various refractive index (Fc_1 and Fc_2, n=4).

#### Ligand density evaluation was then performed to qualify the biosensor quantitative potential

(Pol et al., 2016). Fetuin glycoprotein was covalently grafted on both the sensing and reference surfaces to reach full surface coverage. An enzymatic lysis of terminal sialic acid moieties of the reference surface was achieved by bacterial neuraminidase (Durous et al., 2019). The signal decrease (ΔRU) during the lysis of reference surface (see **Supplementary FIG 4**) was converted into RU/kDa and molecules.mm^-2^. This conversion used the mass sensitivity of SPR systems (1 RIU ↔ 10^-6^ g.mm^-2^), our biosensor calibration and the known molecular weights of the fetuin (48.4 kDa) and sialic acid moieties (309 Da) (Pol et al., 2016; Stenberg, 1990).

**Supplementary FIG4:**
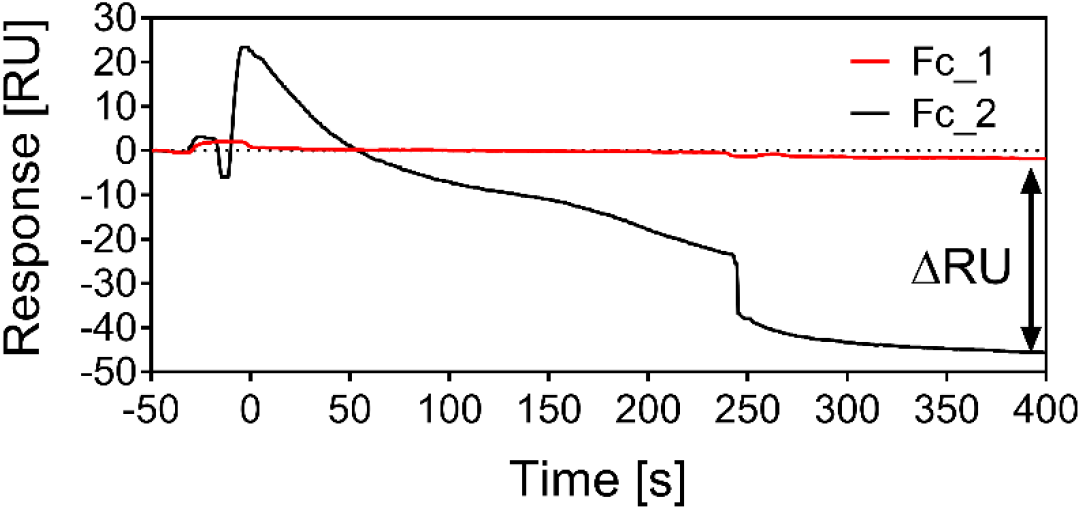
Signal decrease (ΔRU) obtained during the enzymatic lysis with bacterial neuraminidase of reference surface (Fc_2)

The value obtained of **168±17RU.kDa^-1^** fetuin glycoprotein covalently grafted referred to a high surface density of sialic acid terminal moieties on the biosensor surface (see **Supplementary Table 1**). This was described as particularly suited for quantitative SPR analysis (Pol et al., 2016). Additionally, we evaluated that the fetuin glycoproteins covalently immobilized expose an average number of terminal sialic acid (SA) moieties of ***n=3.2*** SA/protein for the binding of influenza virus.

**Supplementary Table 1:**
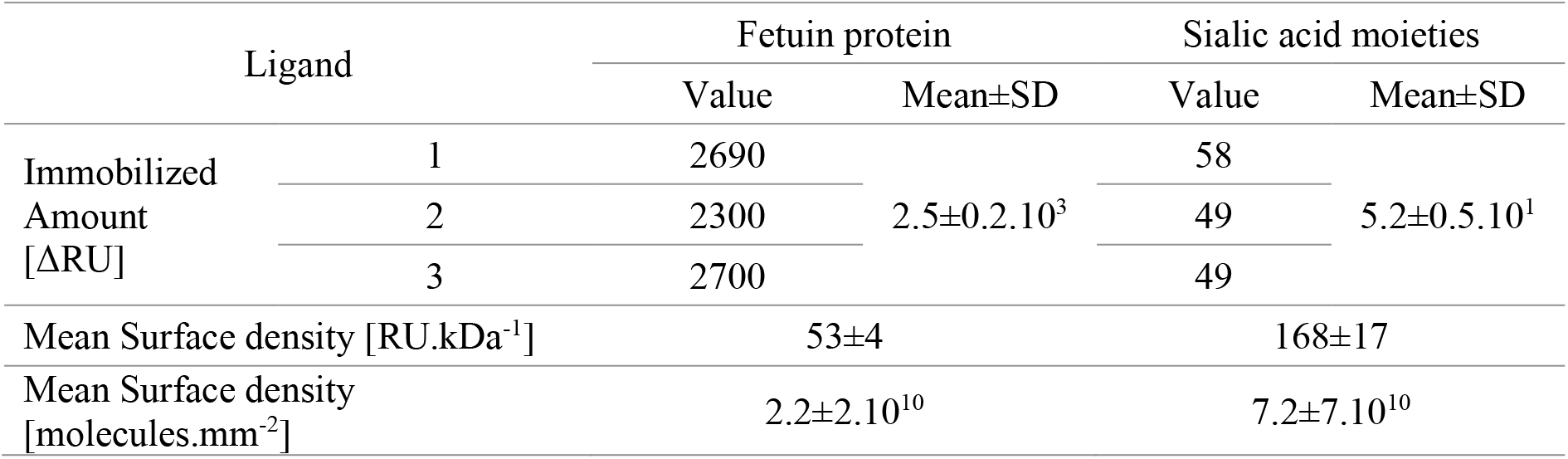
Evaluation of the ligand surface density according to the amount of fetuin immobilized on the reference surface and the signal decrease evaluated during enzymatic lysis (n=3 experiments).

**Supplementary FIG S1 :**
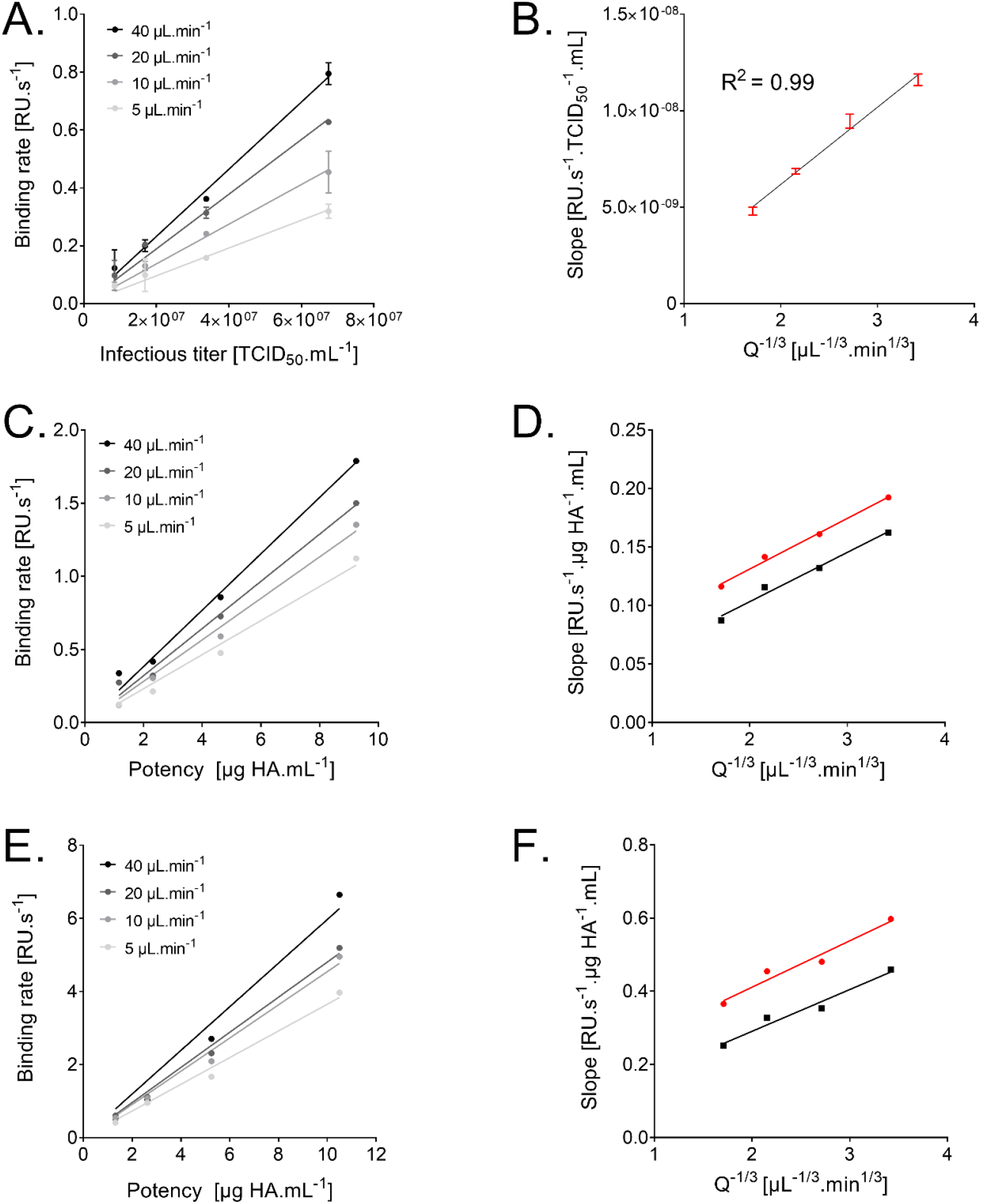
Calibration curves obtained for various influenza virus samples.

### SUPPLEMENTARY INFORMATION 3 Evaluation of influenza virus stability

**Supplementary FIG6.**
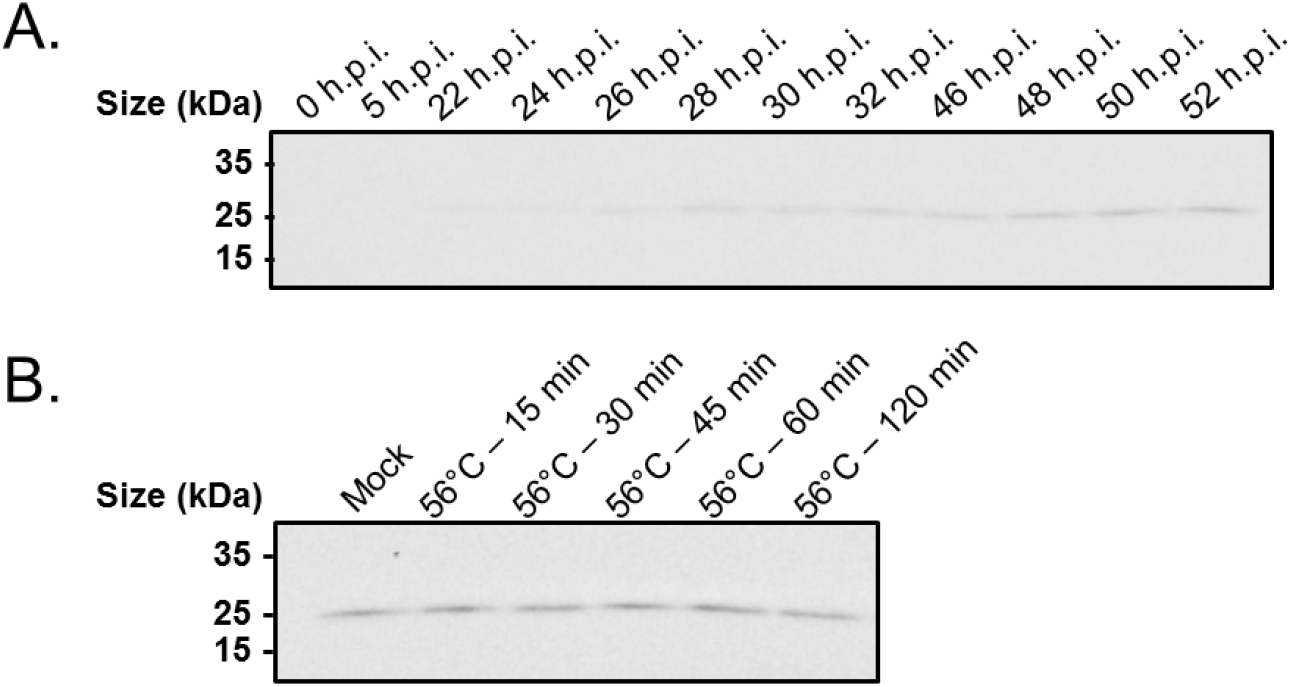
Influenza virus production kinetics and viral particle stability by Western Blot densitometry analysis with universal anti-HA antibodies. A_A/Panama/07/99 (H3N2) production kinetics in DuckCell-T17 suspension cell culture according to time post-infection B_Relative HA content of heat-stressed samples (H3N2 A/Panama/07/99 – 54 hpi) according to sample incubation time at 56°C.

**Supplementary FIG7:**
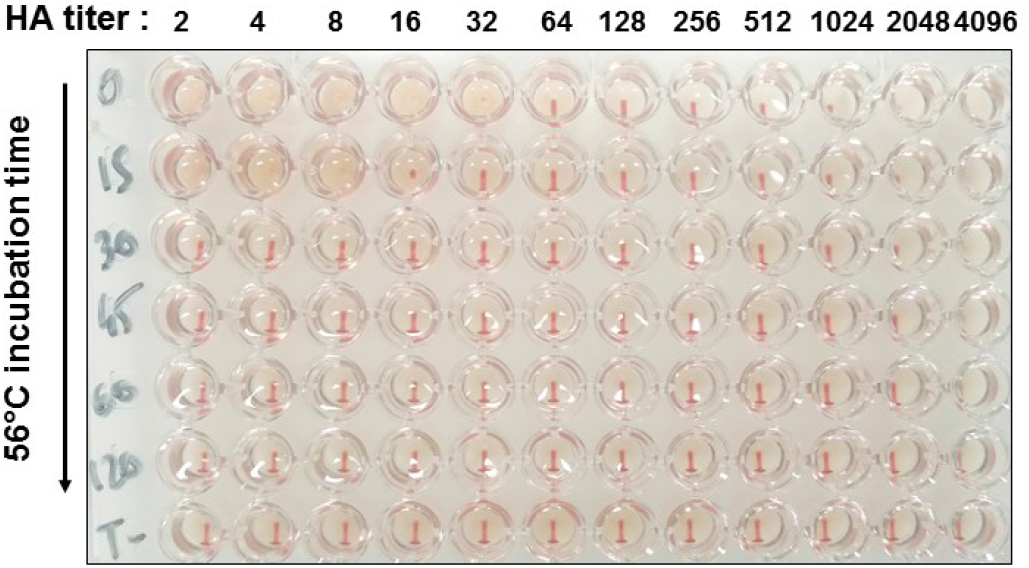
Influenza viral particle stability evaluated by Hemaglutination assay (HA titers). HA titers of H3N2 A/Panama/07/99 produced in DuckCell-T17 suspension cell culture supernatant (54 hpi) were treated for 0 to 120-min at 56°C. Only 0- and 15-min incubation time demonstrated bioactive hemagglutinin evaluated as 32 and 16 HA units/50 μL, respectively.

### SUPPLEMENTARY INFORMATION 4 Description of online SPRi biosensor setup

#### Script for the Arduino microcontroller

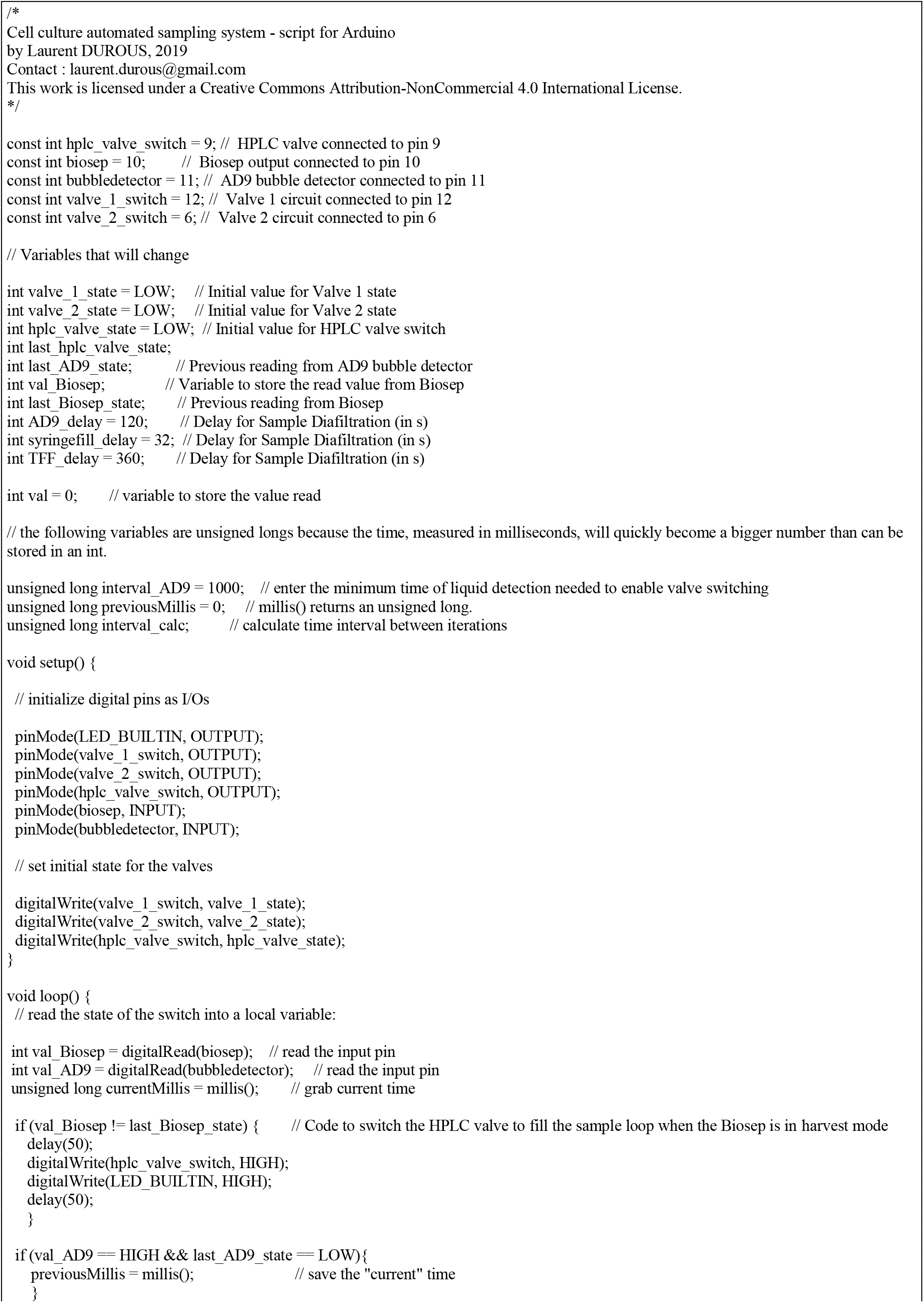
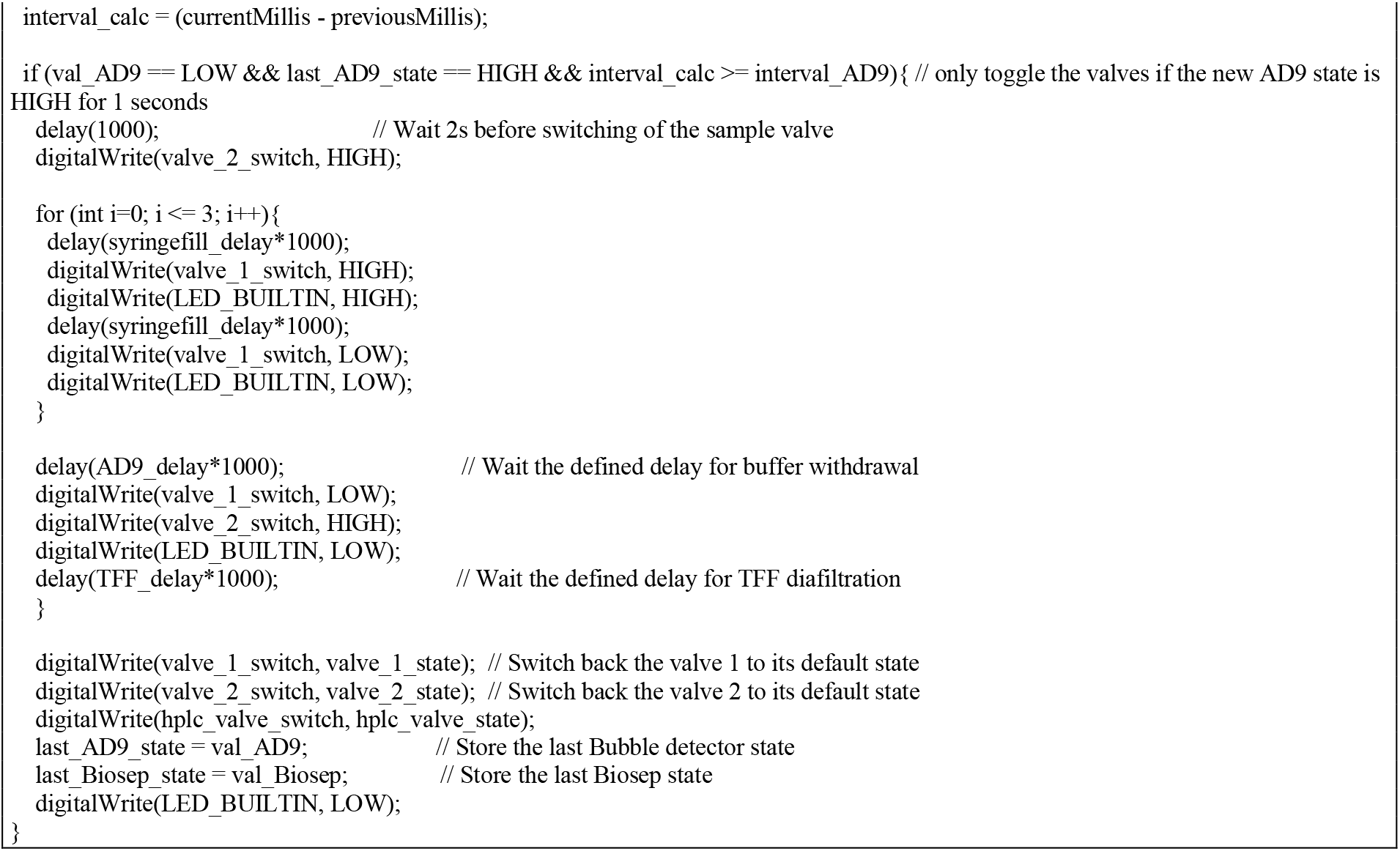

**Supplementary Table 5.**
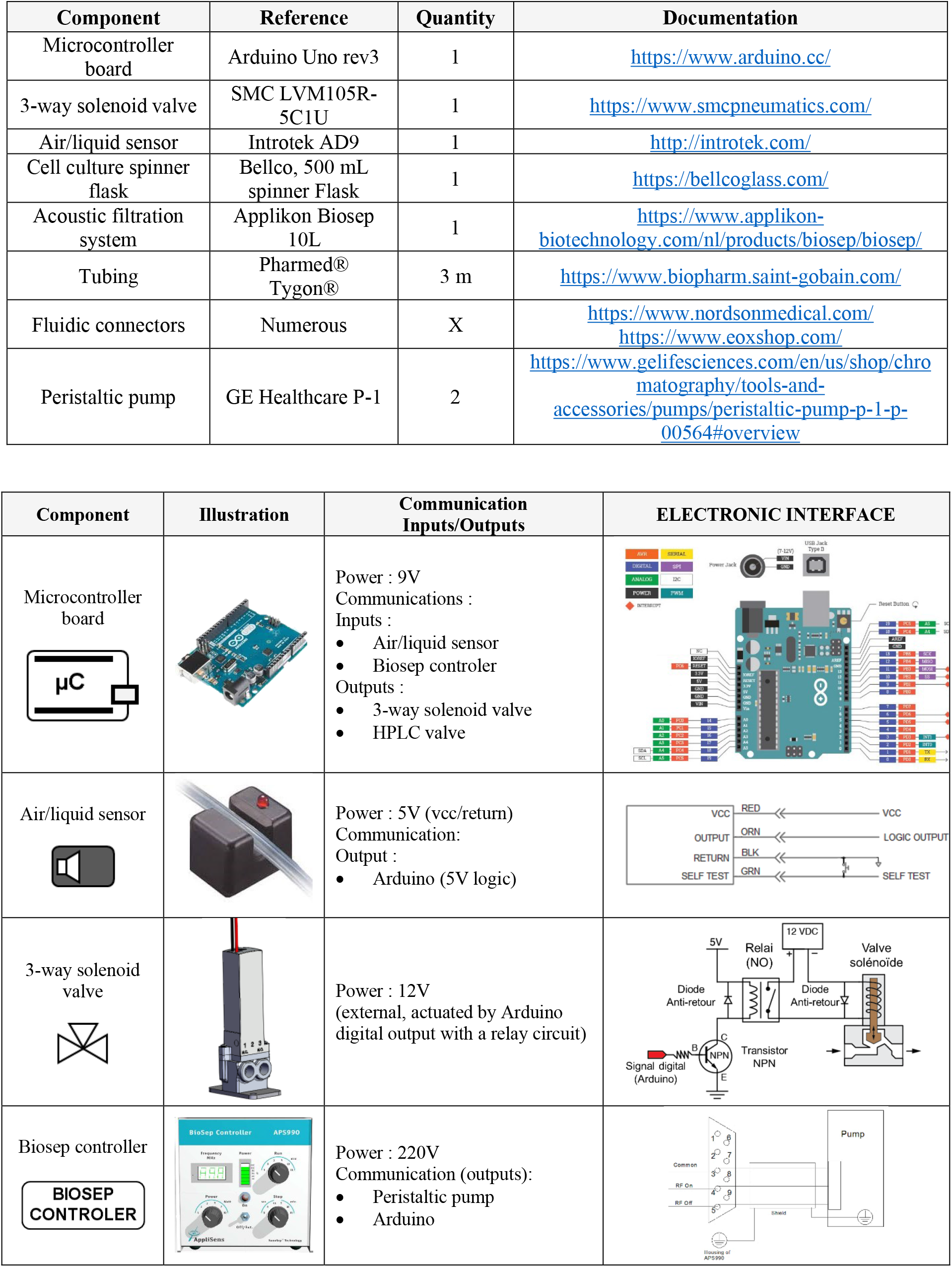
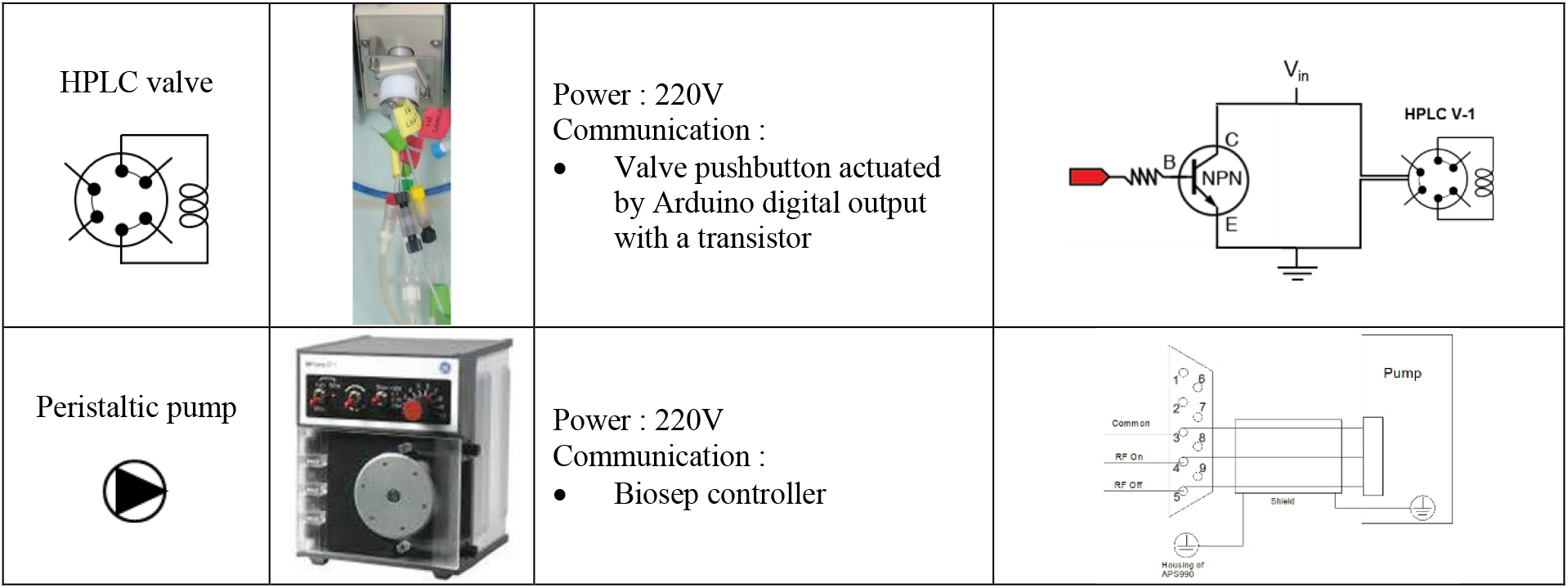
Electronic connections/communications for the automatic sampling and purification line components. Electronic interfaces are based on reference manuals of the components (see ref (Applikon Biotechnology, 2018; GE Healthcare, 2005; Introtek, 2016)).

