## Supplementary Informations for "REAL-TIME AT-LINE MONITORING OF INFLUENZA VIRUS IN CELL CULTURE BY A SURFACE PLASMON RESONANCE BIOSENSOR"

**SUPPLEMENTARY FIG 1 – Impact of virus size variation on the mass transport coefficient $k_{t}$:** Schematic representation of the transport of virus particles in the biosensor flow cell.

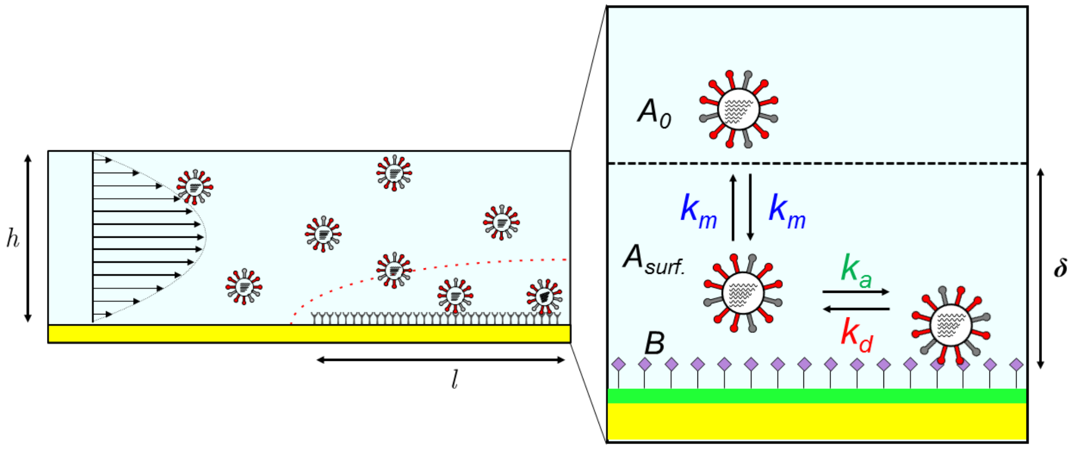


**Supplementary FIG1**_Schematic representation of the transport of virus particles in the biosensor flow cell.

The diffusive flux $J\ [mol.m^{2}.s^{-1}]$ of an analyte through the depletion zone ($\delta$) of a biosensor is given by (Sjölander and Urbaniczky, 1991; Squires et al., 2008):

$$J = k_m\left(A_{bulk} - A_{surf.}\right).10^3 \quad [\boldsymbol{equation\ S1}]$$

Where $k_m\ [m.s^{-1}]$ is the heterogeneous mass transport coefficient and $A_{bulk}$ and $A_{surf.}\ [mol.L^{-1}]$ correspond respectively to the analyte concentrations in the solution and close to the sensor's surface. Factor $10^3$ is used as a conversion factor: $mol.m^{-3} \leftrightarrow mol.L^{-1}$. For a rectangular flow cell with one sensing wall, the heterogeneous mass transport coefficient has been analytically derived (Karlsson, 2016; Pol et al., 2016) :

$$k_m = \ 0{,}98 \times \left[\frac{D^2\ Q}{h^2\ w\ l}\right]^{\frac{1}{3}} \quad [\boldsymbol{equation\ S2}]$$

Where $D$ [m².s] is the diffusion coefficient of the analyte, Q [m³.s⁻¹] is the volumetric flow rate, and h, w, and l [m] respectively correspond to the height, the width and the length of the flow cell. Surface-based biosensors can be operated in conditions of mass transport limitation, where the binding rate of biomolecules to the biosensor's surface is limited by the diffusion phenomenon, predominating over the association phenomenon. Under these conditions, the bulk concentration of the analyte is proportional to the initial binding rate dΓ/dt according to the relation:

$$\left.\frac{d\,\Gamma}{dt}\right|_{t\sim0} \sim k_m\,A_{bulk} \quad [\boldsymbol{equation\ S3}]$$

$\Gamma\ [mol.m^{2}]$ being the surface-bound density of the analyte. The corresponding response of SPR biosensors (e.g., the initial binding rate in terms of response unit) also takes other parameters into account:

$$\left.\frac{dR}{dt}\right|_{t\sim0} = k_t \times A_{bulk} = \ Slope_{cal} \times G\ \times MW_{virus} \times k_m\ \times A_{bulk} \quad [\boldsymbol{equation\ S4}]$$

$Slope_{cal}$ is a calibration coefficient between the SPR response and the local refractive index variation, G is a conversion factor between the refractive index variation and the mass surface density, and $MW_{virus}$ is the molecular weight of Influenza virus particle (Pol et al., 2016).

Besides its flow rate dependence ($k_t \sim Q^{1/3}$), $k_t$ depends on constants (i.e. geometric flow cell parameters and conversion factors) and analyte-related parameters ($D_{virus}$ and $MW_{virus}$).

The size-dependence of the diffusion coefficient of the virus can be evaluated according to the Stokes−Einstein equation:

$$D_{virus} = \frac{k_B \times T}{3\pi \times \eta \times d_{virus}} \ [\boldsymbol{equation\ S5}]$$

Where $k_B$ represents the Boltzmann constant, $T$ the temperature and $\eta$ the viscosity of the medium. The flow cell temperature was kept constant during all experiments. Buffer viscosity was also considered constant as all samples were diluted in the same buffer for calibrations (PBST added with 10µM oseltamivir carboxylate). Additionally, the viscosity of the cell culture medium (Optipro®) was considered identical to PBST buffer, as no serum was added.

While for the molecular weight of influenza virus:

$$MW_{virus} = \frac{\rho_{virus} \times V_{virus}}{N_A} = \frac{\rho_{virus}}{N_A} \times \frac{1}{6}\pi\, d_{virus}^3 \ [\boldsymbol{equation\ S6}]$$

Where $\rho_{virus}$ represents the density and $V_{virus}$ the volume of the virus particle and

By coupling **equations S4, S5 and S6**, we can deduce the mass transport coefficient is proportional to the virus particle diameter according to $\boldsymbol{k_t \sim d_{virus}^{7/3}}$.

**SUPPLEMENTARY INFORMATION 2 – Qualification of the SPRi biosensor and selection of mass transport limitation conditions for the quantitative analysis of influenza virus particles.**

**Calibration of the SPR-2 apparatus** was realized by analyzing glycerol solutions diluted in PBST buffer onto the functionalized sensor. The solutions were of increasing refractive index ranging from 0 to 0.0015 [ΔRIU]. Considering a 1% w/w glycerol solution yields a refractive index increment $\Delta n_s$ of 0.00113 Refractive Index Unit [RIU], we evaluated the system response as:

$$\boldsymbol{Slope_{cal}} = \frac{\boldsymbol{\Delta R}}{\boldsymbol{\Delta n_s}} = \mathbf{7.08 \pm 0.08\,.\,10^5}\ [\boldsymbol{RU.RIU^{-1}}]$$

This value is close to the conventional value of $\mathbf{1.0.10^6}$ $[\boldsymbol{RU.RIU^{-1}}]$ reported in the literature for commercial SPR systems (Tudos and Schasfoort, 2008). The baseline noise of the response was evaluated as σ (R) = 0.1 [RU]. The resolution of the system can be evaluated by the limit of detection $S = 3.3^* \sigma / Slope_{cal} = 5.10^{-7}$ [RIU].

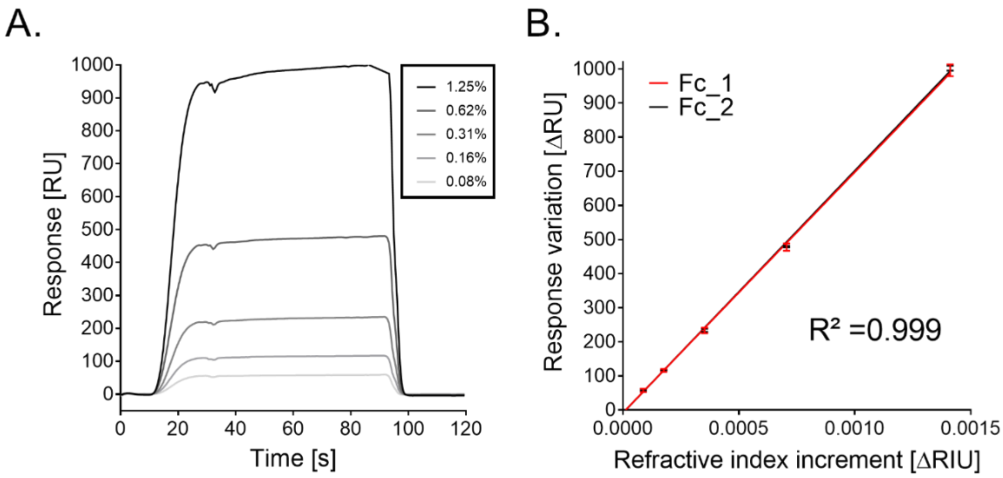

**Supplementary FIG3**: Calibration of the SPR-2 apparatus. A_Sensorgrams of the response for glycerol solutions of increasing concentrations in PBST buffer. B_Calibration curves obtained by difference between response of the reference surface and the sensing surface at the various refractive index (Fc_1 and Fc_2, n=4).

**Ligand density evaluation was then performed to qualify the biosensor quantitative potential** (Pol et al., 2016). Fetuin glycoprotein was covalently grafted on both the sensing and reference surfaces to reach full surface coverage. An enzymatic lysis of terminal sialic acid moieties of the reference surface was achieved by bacterial neuraminidase (Durous et al., 2019). The signal decrease (ΔRU) during the lysis of reference surface (see **Supplementary FIG 4**) was converted into RU/kDa and molecules.$mm^{-2}$. This conversion used the mass sensitivity of SPR systems (1 RIU ↔ $10^{-6}$ g.$mm^{-2}$), our biosensor calibration and the known molecular weights of the fetuin (48.4 kDa) and sialic acid moieties (309 Da) (Pol et al., 2016; Stenberg, 1990).

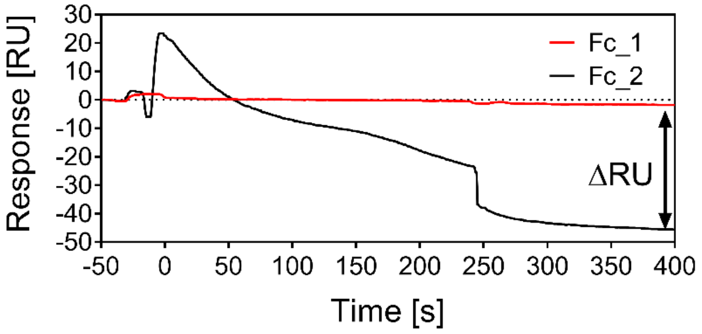

**Supplementary FIG4**: Signal decrease (ΔRU) obtained during the enzymatic lysis with bacterial neuraminidase of reference surface (Fc_2)

The value obtained of **168±17RU.kDa$^{-1}$** fetuin glycoprotein covalently grafted referred to a high surface density of sialic acid terminal moieties on the biosensor surface (see **Supplementary Table 1**). This was described as particularly suited for quantitative SPR analysis (Pol et al., 2016). Additionally, we evaluated that the fetuin glycoproteins covalently immobilized expose an average number of terminal sialic acid (SA) moieties of ***n=3.2*** SA/protein for the binding of influenza virus.

**Supplementary Table 1**: Evaluation of the ligand surface density according to the amount of fetuin immobilized on the reference surface and the signal decrease evaluated during enzymatic lysis (n=3 experiments).

| Ligand | | Fetuin protein | | Sialic acid moieties | |
|---|---|---|---|---|---|
| | | Value | Mean±SD | Value | Mean±SD |
| Immobilized Amount [ΔRU] | 1 | 2690 | $2.5\pm0.2.10^{3}$ | 58 | $5.2\pm0.5.10^{1}$ |
| | 2 | 2300 | | 49 | |
| | 3 | 2700 | | 49 | |
| Mean Surface density [RU.kDa$^{-1}$] | | 53±4 | | 168±17 | |
| Mean Surface density [molecules.mm$^{-2}$] | | $2.2\pm2.10^{10}$ | | $7.2\pm7.10^{10}$ | |

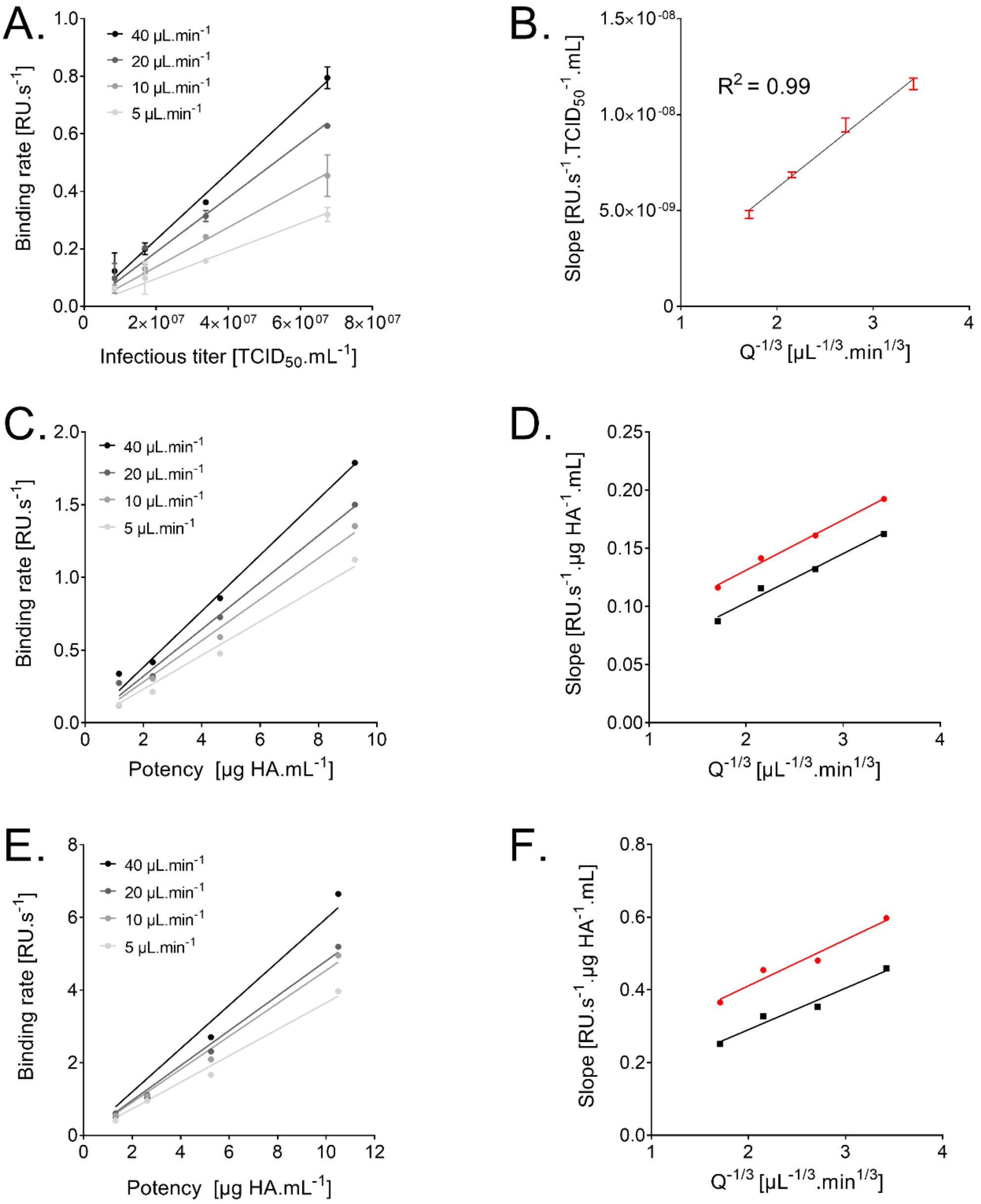


**Supplementary FIG S1** : Calibration curves obtained for various influenza virus samples.

**SUPPLEMENTARY INFORMATION 3 – Evaluation of influenza virus stability.**

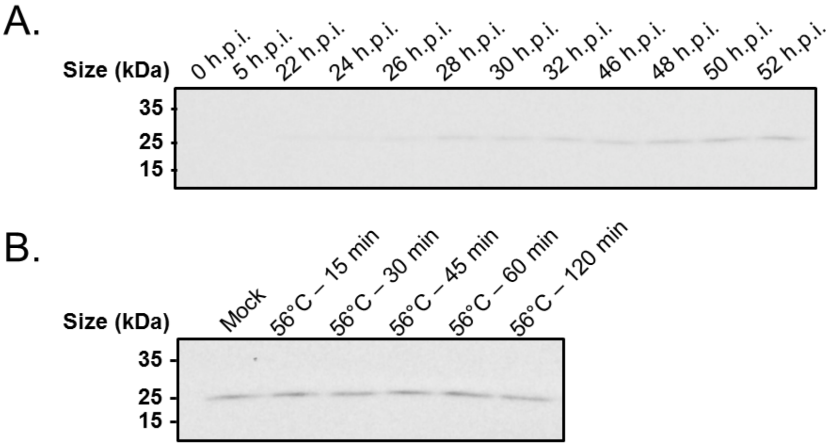


**Supplementary FIG6_ Influenza virus production kinetics and viral particle stability by Western Blot densitometry analysis with universal anti-HA antibodies.** A_A/Panama/07/99 (H3N2) production kinetics in DuckCell-T17 suspension cell culture according to time post-infection B_Relative HA content of heat-stressed samples (H3N2 A/Panama/07/99 – 54 hpi) according to sample incubation time at 56°C.

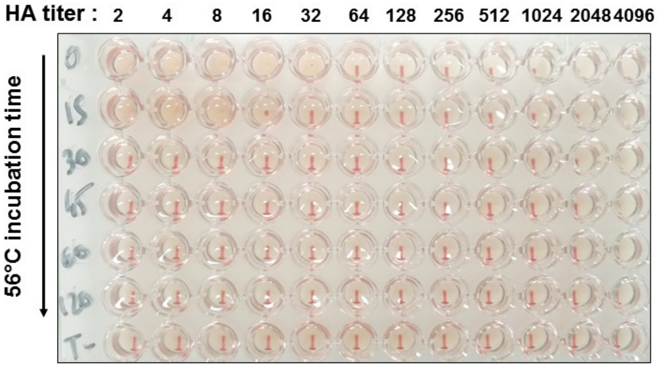


**Supplementary FIG7_ Influenza viral particle stability evaluated** by Hemaglutination assay (HA titers). HA titers of H3N2 A/Panama/07/99 produced in DuckCell-T17 suspension cell culture supernatant (54 hpi) were treated for 0 to 120-min at 56°C. Only 0- and 15-min incubation time demonstrated bioactive hemagglutinin evaluated as 32 and 16 HA units/50 µL, respectively.

### SUPPLEMENTARY INFORMATION 4 – Description of online SPRi biosensor setup.

#### Script for the Arduino microcontroller

```
/*
Cell culture automated sampling system - script for Arduino
by Laurent DUROUS, 2019
Contact :
This work is licensed under a Creative Commons Attribution-NonCommercial 4.0 International License.
*/

const int hplc_valve_switch = 9; //  HPLC valve connected to pin 9
const int biosep = 10;         //  Biosep output connected to pin 10
const int bubbledetector = 11; //  AD9 bubble detector connected to pin 11
const int valve_1_switch = 12; //  Valve 1 circuit connected to pin 12
const int valve_2_switch = 6; //  Valve 2 circuit connected to pin 6

// Variables that will change

int valve_1_state = LOW;     // Initial value for Valve 1 state
int valve_2_state = LOW;     // Initial value for Valve 2 state
int hplc_valve_state = LOW;  // Initial value for HPLC valve switch
int last_hplc_valve_state;
int last_AD9_state;           // Previous reading from AD9 bubble detector
int val_Biosep;              // Variable to store the read value from Biosep
int last_Biosep_state;        // Previous reading from Biosep
int AD9_delay = 120;        // Delay for Sample Diafiltration (in s)
int syringefill_delay = 32;  // Delay for Sample Diafiltration (in s)
int TFF_delay = 360;        // Delay for Sample Diafiltration (in s)

int val = 0;        // variable to store the value read

// the following variables are unsigned longs because the time, measured in milliseconds, will quickly become a bigger number than can be
stored in an int.

unsigned long interval_AD9 = 1000;    // enter the minimum time of liquid detection needed to enable valve switching
unsigned long previousMillis = 0;     // millis() returns an unsigned long.
unsigned long interval_calc;          // calculate time interval between iterations

void setup() {

  // initialize digital pins as I/Os

  pinMode(LED_BUILTIN, OUTPUT);
  pinMode(valve_1_switch, OUTPUT);
  pinMode(valve_2_switch, OUTPUT);
  pinMode(hplc_valve_switch, OUTPUT);
  pinMode(biosep, INPUT);
  pinMode(bubbledetector, INPUT);

  // set initial state for the valves

  digitalWrite(valve_1_switch, valve_1_state);
  digitalWrite(valve_2_switch, valve_2_state);
  digitalWrite(hplc_valve_switch, hplc_valve_state);
}

void loop() {
  // read the state of the switch into a local variable:

 int val_Biosep = digitalRead(biosep);    // read the input pin
 int val_AD9 = digitalRead(bubbledetector);     // read the input pin
 unsigned long currentMillis = millis();        // grab current time

  if (val_Biosep != last_Biosep_state) {        // Code to switch the HPLC valve to fill the sample loop when the Biosep is in harvest mode
     delay(50);
     digitalWrite(hplc_valve_switch, HIGH);
     digitalWrite(LED_BUILTIN, HIGH);
     delay(50);
     }

  if (val_AD9 == HIGH && last_AD9_state == LOW){
      previousMillis = millis();                         // save the "current" time
      }
```

```
  interval_calc = (currentMillis - previousMillis);

  if (val_AD9 == LOW && last_AD9_state == HIGH && interval_calc >= interval_AD9){ // only toggle the valves if the new AD9 state is HIGH for 1 seconds
    delay(1000);                                    // Wait 2s before switching of the sample valve
    digitalWrite(valve_2_switch, HIGH);

    for (int i=0; i <= 3; i++){
      delay(syringefill_delay*1000);
      digitalWrite(valve_1_switch, HIGH);
      digitalWrite(LED_BUILTIN, HIGH);
      delay(syringefill_delay*1000);
      digitalWrite(valve_1_switch, LOW);
      digitalWrite(LED_BUILTIN, LOW);
    }

    delay(AD9_delay*1000);                          // Wait the defined delay for buffer withdrawal
    digitalWrite(valve_1_switch, LOW);
    digitalWrite(valve_2_switch, HIGH);
    digitalWrite(LED_BUILTIN, LOW);
    delay(TFF_delay*1000);                          // Wait the defined delay for TFF diafiltration
    }

    digitalWrite(valve_1_switch, valve_1_state);  // Switch back the valve 1 to its default state
    digitalWrite(valve_2_switch, valve_2_state);  // Switch back the valve 2 to its default state
    digitalWrite(hplc_valve_switch, hplc_valve_state);
    last_AD9_state = val_AD9;                       // Store the last Bubble detector state
    last_Biosep_state = val_Biosep;                 // Store the last Biosep state
    digitalWrite(LED_BUILTIN, LOW);
}
```

**Supplementary Table 5** _ Electronic connections/communications for the automatic sampling and purification line components. Electronic interfaces are based on reference manuals of the components (see ref (Applikon Biotechnology, 2018; GE Healthcare, 2005; Introtek, 2016)).

| Component | Reference | Quantity | Documentation |
|---|---|---|---|
| Microcontroller board | Arduino Uno rev3 | 1 | https://www.arduino.cc/ |
| 3-way solenoid valve | SMC LVM105R-5C1U | 1 | https://www.smcpneumatics.com/ |
| Air/liquid sensor | Introtek AD9 | 1 | http://introtek.com/ |
| Cell culture spinner flask | Bellco, 500 mL spinner Flask | 1 | https://bellcoglass.com/ |
| Acoustic filtration system | Applikon Biosep 10L | 1 | https://www.applikon-biotechnology.com/nl/products/biosep/biosep/ |
| Tubing | Pharmed® Tygon® | 3 m | https://www.biopharm.saint-gobain.com/ |
| Fluidic connectors | Numerous | X | https://www.nordsonmedical.com/ https://www.eoxshop.com/ |
| Peristaltic pump | GE Healthcare P-1 | 2 | https://www.gelifesciences.com/en/us/shop/chromatography/tools-and-accessories/pumps/peristaltic-pump-p-1-p-00564#overview |

| Component | Illustration | Communication Inputs/Outputs | ELECTRONIC INTERFACE |
|---|---|---|---|
| Microcontroller board 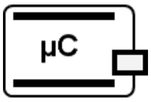 | 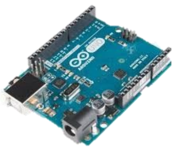 | Power : 9V<br>Communications :<br>Inputs :<br>• Air/liquid sensor<br>• Biosep controler<br>Outputs :<br>• 3-way solenoid valve<br>• HPLC valve | 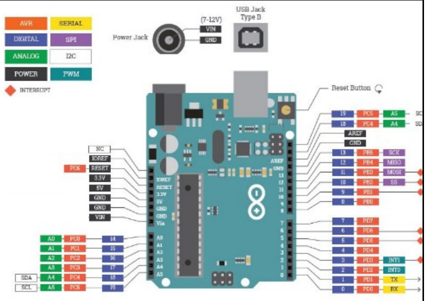 |
| Air/liquid sensor 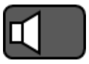 | 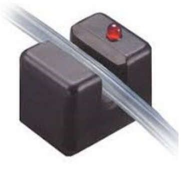 | Power : 5V (vcc/return)<br>Communication:<br>Output :<br>• Arduino (5V logic) | 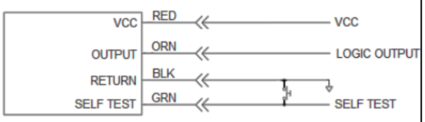  |
| 3-way solenoid valve 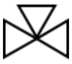 | 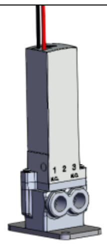 | Power : 12V<br>(external, actuated by Arduino digital output with a relay circuit) | 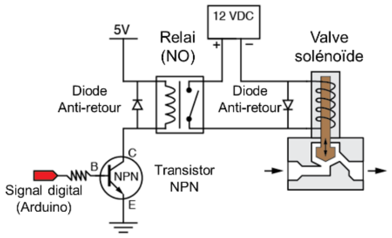  |
| Biosep controller 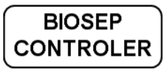  | 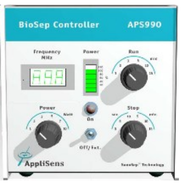  | Power : 220V<br>Communication (outputs):<br>• Peristaltic pump<br>• Arduino | 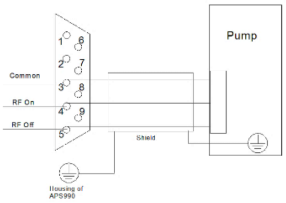  |

| | | | |
|---|---|---|---|
| HPLC valve 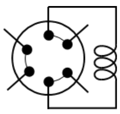 | 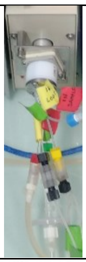 | Power : 220V<br>Communication :<br>• Valve pushbutton actuated by Arduino digital output with a transistor | 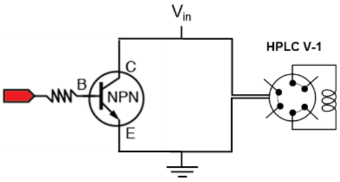 |
| Peristaltic pump 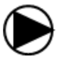 | 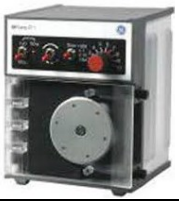 | Power : 220V<br>Communication :<br>• Biosep controller | 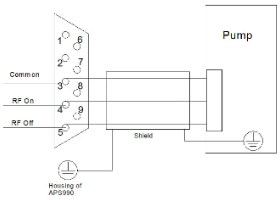 |
